# Non-canonical Wnt signalling initiates scarring in biliary disease

**DOI:** 10.1101/276196

**Authors:** DH Wilson, RP Mellin, NT Younger, EJ Jarman, A Raven, P Chen, CH Dean, DJ Henderson, TJ Kendall, L Boulter

**Author notes:** Corresponding Author: Dr Luke Boulter. MRC Human Genetics Unit, Institute of Genetic and Molecular Medicine, Edinburgh, EH4 2XU.

## Abstract

Cholangiopathies, or biliary diseases, account for a significant proportion of adult and paediatric liver disease. In these pathologies, iterative cycles of damage and repair result in the development of a regenerative microenvironment surrounding the bile ducts, which orchestrates both epithelial proliferation and also biliary fibrosis. Ultimately, fibrosis at the cost of repair results in cholestasis and liver failure, necessitating liver transplantation. Whilst the fibrogenic mechanisms in hepatocellular disease have been widely studied, little is known about the processes that regulate biliary scarring. We sought to determine how the injured biliary epithelium communicates to adjacent stromal cells to regulate scar formation, and to identify therapeutically targetable pathways that could be inhibited to reduce biliary scarring, whilst maintaining the pro-regenerative stroma. Using human tissue, bile duct organoids and animal models of biliary disease, we show that non-canonical Wnt signalling is important in initiating biliary scarring. This process is driven by myeloid Wnt5a and acts through epithelial Vangl2, which is upstream of Jnk/cJun signalling. Activation of this pathway drives a pro-fibrotic signalling process which instructs portal fibroblasts to synthesise collagen. Finally, we determine that therapeutic Wnt ligand inhibition reduces biliary scarring, identifying non-canonical Wnt signalling as a novel target for anti-fibrotic therapy in cholestatic biliary disease.

## Introduction

Biliary diseases, also known as cholangiopathies, account for approximately one-third of all adult liver disease and 70% of childhood liver disease. However, compared to their parenchymal counterparts, where hepatoyctes are damaged, the mechanisms that drive the progression of cholangiopathies are unknown and therapeutics to target the formation of scar tissue have not been forthcoming(1). As a result, for the majority of patients with chronic biliary disease, transplantation is the only curative option(2).

At the tissue level, biliary disease and regeneration relies on linking the proliferation of biliary epithelial cells (cholangiocytes) with the formation of a regenerative microenvironment that provides both inductive cues to the epithelium but also maintains tissue integrity through scar deposition(3, 4). How these different cellular components are integrated to form a regenerative hub is unknown.

In many contexts, epithelial regeneration is controlled through activation of the canonical Wnt signalling pathway, which relies on the translocation of β-catenin into the nucleus to drive changes in gene expression(5–9). In reality, Wnt ligands are integrated into a much larger network, activating a number of Wnt dependant signalling cascades that function independent of β-catenin(10–12). These so called non-canonical Wnt signalling pathways are highly conserved from *Drosophila*(13) through to vertebrates(14) and utilise Wnt ligands in combination with a broad range of cell surface receptors, to activate diverse downstream processes(15). The most widely reported non-canonical Wnt signalling cascades are planar cell polarity (PCP), which regulates activation of cJUN dependant transcription and cytoskeletal rearrangement through Rho and Rock(16–18), and Wnt/Ca^2+^ signalling, which, though modulation of the cellular Ca^2+^ concentration, activates NFAT dependant transcription(19–21). Other non-canonical Wnt signalling pathways have also been identified; for example activation of YAP/TAZ signalling though Rho dependant inhibition of Lats1/2, however it remains unclear how broadly applicable these pathways are(22).

A small number of studies have demonstrated that non-canonical Wnt signalling plays a role in post-natal organ growth(23–25) and in cancer(26–28). These studies have shown that, in the adult, non-canonical Wnt components are involved in the activation of cJun N-terminal kinase (JNK)(17), regulation of matrix-metalloproteinase-14(29), activation of integrin αv(30) and promotion of epithelial polarity(25). Moreover, a small number of reports have indicated that non-canonical Wnt signalling, through Vangl2, regulates the extra-cellular matrix *in vivo*(4, 25, 31). However, it remains unclear what the core function of non-canonical Wnt signalling is in the adult liver.

In this study, we demonstrate that a group of Wnt ligands associated with non-canonical Wnt signalling, particularly Wnt4 and Wnt5a, are up regulated rapidly in a model of biliary disease and these ligands are also highly expressed in patients with primary sclerosing cholangitis (a highly fibrotic biliary disease with limited treatment options). Therapeutic inhibition of the Wnt signalling pathway in this context, through the prevention of Wnt ligand secretion, does not affect the proliferation of biliary cells or the number of regenerating bile ducts. It does, however, reduce the level of fibrosis deposited around these proliferating ducts. We go on to demonstrate that myeloid Wnt5a regulates this process by activating JNK/cJUN signalling specifically in biliary epithelial cells, promoting their cross talk with portal fibroblasts thereby regulating the capacity of these cells to synthesise collagen and form scar tissue. This study represents, to our knowledge, the first description of how non-canonical Wnt signalling functions to regulate adult tissue scarring by integrating a number of cell types and offers a novel therapeutic target to treat biliary diseases in patients.

## Results

### The non-canonical Wnt pathway is activated in biliary disease

Previous studies have shown that canonical Wnt signalling is not activated in the injured bile duct (through lack of nuclear accumulation of β-catenin)(5, 32), however, canonical pathway activity occurs as bile ducts become malignant and the canonical Wnt signal drives tumour progression(33, 34). Here we show that in biliary epithelial cells from patients who have undergone transplantation for Primary Sclerosing Cholangitis (PSC), and non-PSC patient livers, β-catenin is detected at the cell membrane but not in the nucleus (figure 1a, left-hand panels), indicating that the canonical Wnt pathway is not active. Furthermore, there was no detectable expression of the canonical Wnt targets LEF1 and cMYC in these cells, in both uninjured and PSC cases (figure 1a, centre and right-hand panels). Activation of non-canonical Wnt signalling, through pathways that are linked to PCP signalling, can be determined through staining for activation of JNK and its substrate, cJUN. In biliary cells of uninjured livers, there is a low level of pJNK^T183/Y185^, cJUN and p-cJUN^S73^ in biliary cells (figure 1b, upper panels), however in PSC, biliary cells are positive for p-JNK^T183/Y185^, and p-cJUN^S73^, indicating that in biliary disease, non-canonical Wnt signalling could contribute to the activation of JNK-cJUN dependant transcription (figure 1b, lower panels).

**Figure 1.**
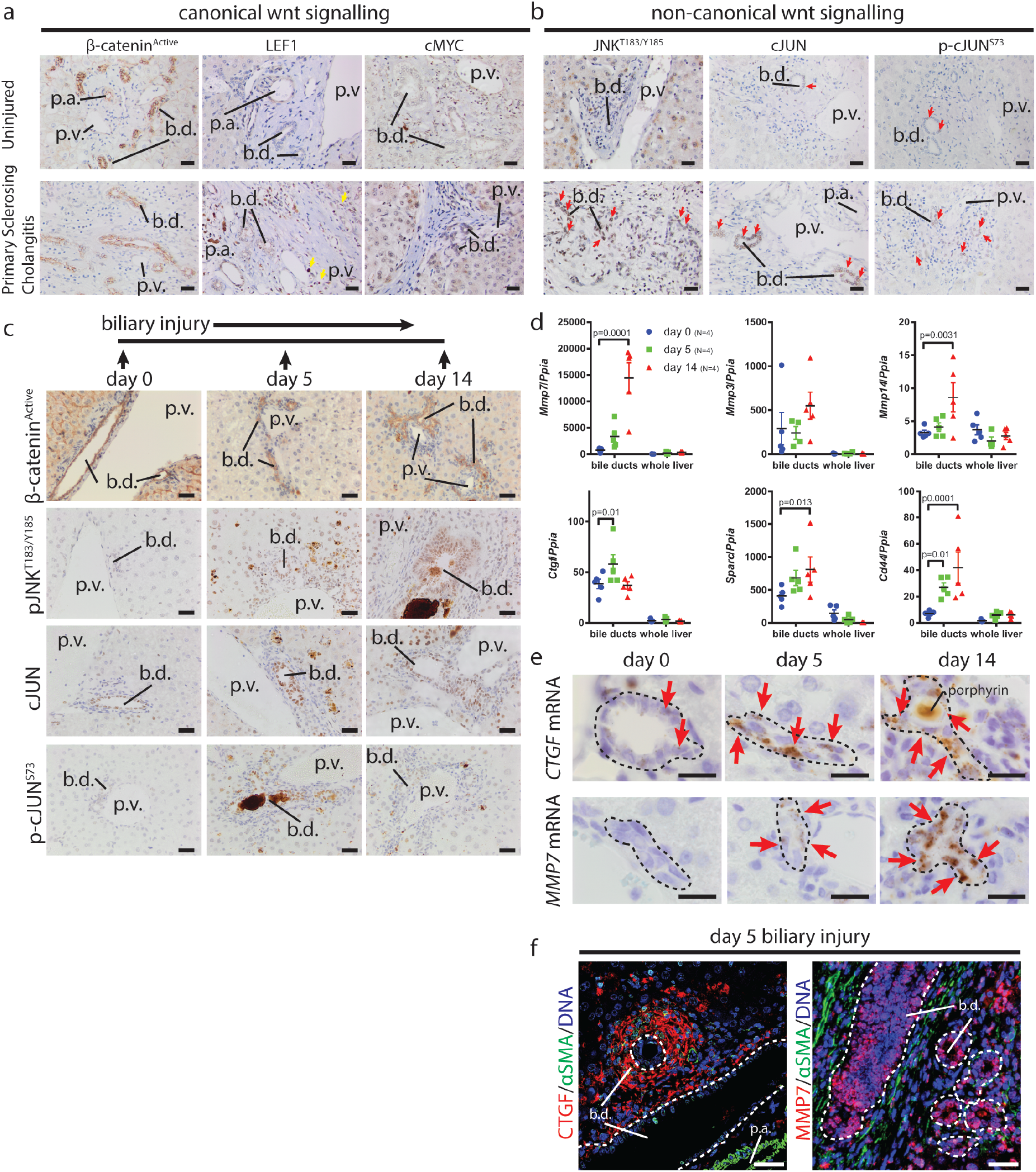
The non-canonical Wnt signalling pathway is activated during cholestatic liver disease. (a) Immunostaining of canonical Wnt signalling intermediates (active β-catenin, Lef1 and cMyc) (b) Immunostaining of non-canonical Wnt signalling intermediates (p-JNK1/2^T183/Y185^, cJUN and p-cJUN^S73^) in uninjured liver and primary sclerosing cholangitis. (c) Schematic representation of the DDC model with staining of active β-catenin, p-Jnk^T183/Y185^, cJUN and p-cJUN^S73^ at 0, 5 and 14 days following dietary induction. (d) Relative mRNA expression of Wnt signalling targets *Mmp3*, *Mmp7*, *Mmp14*, *Cd44*, *Sparc*, *Ctgf* in isolated bile ducts and whole liver thought-out the DDC time course. (e) RNAscope for *Ctgf* and *Mmp7* at day 0, day 5 and day 14 following DDC injury, red arrows denote positivity. (f) Immunofluorescence staining of day 5 DDC liver for CTGF or MMP7 (red) and Fibroblasts, (αSMA, green). Data are presented as mean ±SEM, statistics presented are derived from a Kruskal-Wallis test. Photomicrograph scale bars: 50μm. b.d: bile ducts, p.v. portal vein.

Primary Sclerosing Cholangitis is diagnosed once it is established and as such it is difficult to ascertain whether JNK-cJUN signalling is activated in response to injury and whether the lack of nuclear β-catenin staining is due to this tissue coming from end-stage disease. The 3,5-diethoxycarbonyl-1,4-dihydrocollidine (DDC) model of biliary injury recapitulates many of the features of PSC(35), with ductular occlusions, biliary injury, ductular proliferation and fibrosis. In a time-course of DDC induced biliary injury, induced by administration of 0, 5 or 14 days dietary DDC, proliferative, regenerating, biliary cells were positive for β-catenin, however, as previously shown, β-catenin was restricted to the plasma membrane (figure 1c, upper panels). Furthermore, using a TCF/LEF-H2B:GFP transgenic mouse, which reports activation of canonical Wnt signalling at a single cell resolution(36), we were unable to detect increases in GFP fluorescence with flow cytometry or immunocytochemistry (supplementary figure 1a and 1b) between untreated or day 5 DDC treated mice, nor were we able to detect gene expressions changes of the Wnt signalling pathway targets *Lef1*, *Myc*, *Axin2* or *Ccnd1* in isolated bile ducts, indicating low canonical Wnt pathway activity in enriched biliary cells, which is not upregulated early in injury (supplementary figure 1c-d). Throughout the DDC time course, there is, however, a progressive increase in pJNK^T183/Y185^ and whilst JUN could be detected in the bile ducts at all time points, pJUN^S73^ was only detectable at day 5 following DDC initiation (figure 1c, lower panels). By day 14, pJUN^S73^ was less obvious in the nucleus of biliary cells, suggesting a burst of JUN phosphorylation followed by a tonic level of pathway activation thereafter. In line with activation of JNK/cJUN signalling, expression of cJUN targets, *Mmp7*, *Mmp3*, *Mmp14*, *Cd44* and *Sparc* were increased sequentially over the time course of the study (figure 1d), specifically in bile ducts but not in whole liver tissue. Some of these targets are known canonical pathway targets as well as targets of other transcription factors, so we cannot rule out signalling contributions from β-catenin. However, as β-catenin fails to translocate to the nucleus we suggest that in this context, their transcription is driven by other means. *Ctgf* expression increased transiently in bile ducts, peaking at day 5 following injury, however mRNA levels returned to baseline by day 14 (figure 1d). The expression of a number of potential non-canonical Wnt targets including *Tpm*, *Stat6*, *Rybp*, *Egfr* and *Pkcθ* do not change during the time course of injury and several, *Dcn1*, *Gas1*, *Dkk1*, *Cyr61*, *Bmp4* and *Wnt5b* are decreased over the time course explored (supplementary figure 1e-g). Mmp7 and Ctgf levels have both been implicated in cholestatic disease progression and fibrosis(37–40), however, it remains unresolved whether biliary epithelial cells or portal fibroblasts produce pro-fibrogenic Ctgf and whether companion MMPs regulate the spatial activation of CTGF (41). Using RNAScope, *Ctgf* and *Mmp7* transcripts are rarely seen in the resting liver, however, throughout the DDC time-course, *Ctgf* and *Mmp7* are localised to regenerating biliary epithelial cells (figure 1e), confirming that, in this model, they and not portal fibroblasts are responsible for producing the majority of *Ctgf* and *Mmp7.* Despite this epithelial origin, CTGF localises to adjacent αSMA-positive fibroblasts (figure 1f), suggesting biliary epithelial cells can directly influence their local microenvironment by signalling to adjacent fibrogenic cells. Interestingly, MMP7, which is known to activate CTGF, is restricted to the biliary epithelium potentially representing a localised mechanism by which CTGF is activated.

### Inhibition of Wnt ligand production results in reduced biliary fibrosis

Previous studies have shown that loss of Wnt ligand export in myeloid (using LysM-Cre)(32, 42) or epithelial (using Albumin-Cre)(32) cells through *Wls* deletion, can alter the number of regenerating ductules or levels of fibrosis in biliary disease. However it remains unclear whether this effect is due to altered canonical or non-canonical Wnt signalling. Furthermore, these approaches ablate all Wnt ligands in these lineages in early life and so it remains unresolved as to whether the phenotypes seen are due to abnormal liver maturation in the neonate. To resolve these questions, wild-type mice undergoing DDC induced biliary damage and repair were given LGK974 (from here referred to as Porcupine-i), a potent and highly specific inhibitor of Porcuine(43), which is required to add lipid modifications to Wnt ligands(44). Porcupine-i renders Wnt ligands inactive by preventing ligand export. The benefit of this strategy is that it offers a global, non-lineage specific inhibition of Wnt signalling in the adult liver (figure 2a).

**Figure 2.**
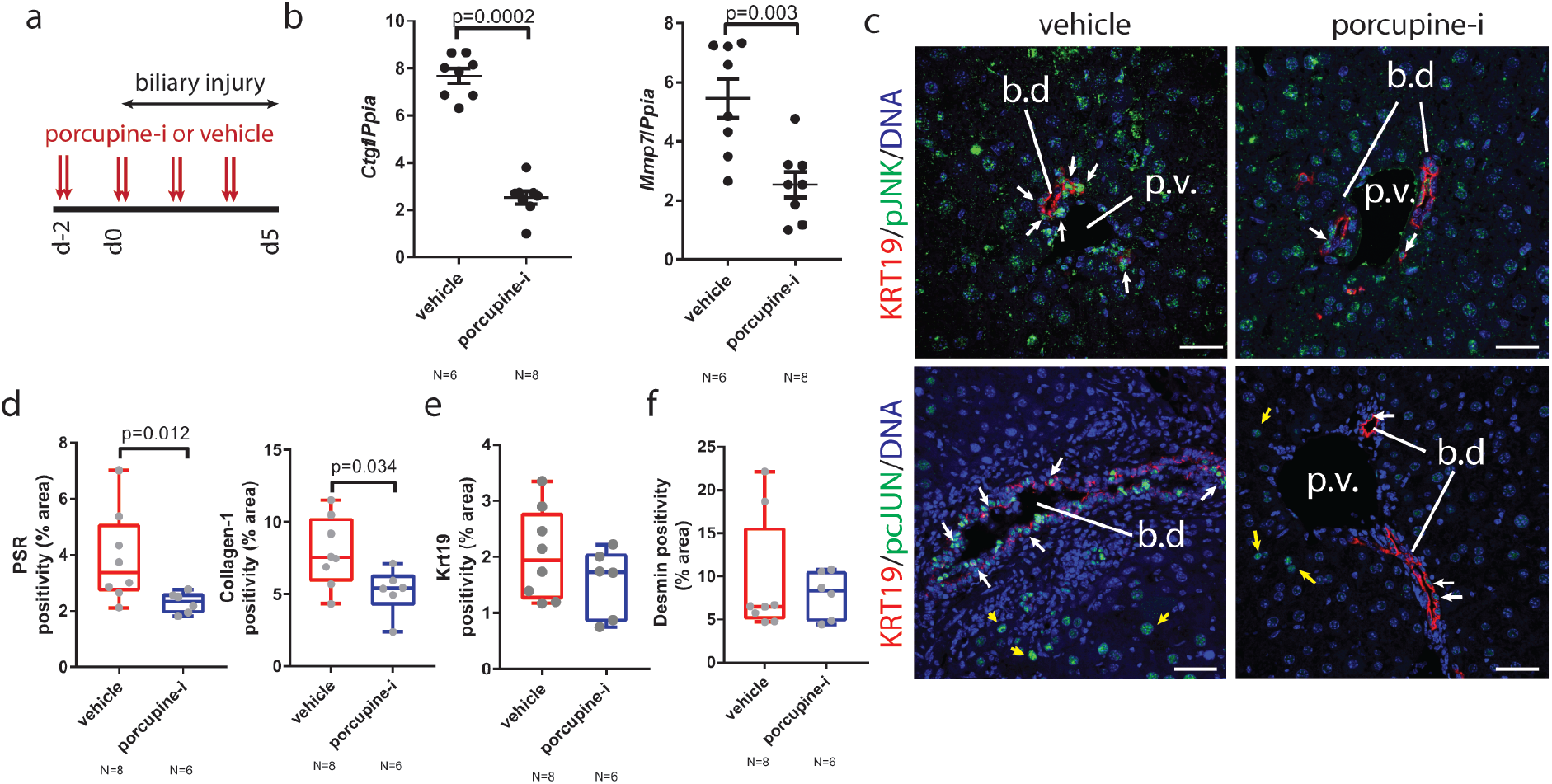
Inhibition of Wnt palmitoylation by Porcupine inhibition results in a reduced biliary fibrosis. (a) Schematic demonstrating dosing regimen during DDC injury of Porcupine inhibitor. (b) Relative *Ctgf* and *Mmp7* mRNA expression in isolated bile ducts from mice treated with Porcupine inhibitor or vehicle. (c) Immunofluorescent staining of bile ducts from mice treated with Porcupine inhibitor or vehicle (Krt19, red) and p-JNK^Thr183/Tyr185^ (upper-panels) and p-cJUN^S73^ (lower-panels), green. White arrows denote positivity. (d) Histological quantification of fibrillar collagen (PSR) and Collagen-1 (e) Keratin-19 and (f) Desmin positive cells in DDC livers from mice treated with Porcupine inhibition or vehicle alone. Data are presented as mean ±SEM, statistics presented are derived from a Mann-Whitney test. Photomicrograph scale bars: 50μm. b.d: bile ducts, p.v. portal vein.

Following administration of Porcupine-i in DDC induced biliary injury; isolated bile ducts had a reduced expression of *Ctgf*, *Mmp7* and *Wnt5b* (figure 2b, supplementary figure 2a), but not *Bmp4*, *Dkk1* or *Cyr61* (supplementary figure 2a). There was also no transcriptional change in levels of Wnt targets *Myc*, *Axin2*, *Lef1* and *Ccnd1* (supplementary figure 2b) or *cJUN* targets *Cd44*, *Sprc*, *Mmp14* and *Mmp3* (supplementary figure 2c). However, there was an obvious reduction in Glutamine Synthetase (GS) positivity in hepatocytes, a known target of canonical Wnt signalling in the liver (supplementary figure 2d), indicating that Wnt signalling had been successfully inhibited by Porcupine-i. In Porcupine-i treated mice which had DDC induced injury, inhibition results in the loss of both pJNK^T183/Y185^ (figure 2c, upper panels) and p-cJUN^S73^ (figure 2c, lower panels) specifically in biliary epithelial cells when compared to DDC animals that received vehicle alone. The consequence of Porcupine inhibition, at the tissue level, is a significant reduction in the levels of fibrillar collagens, determined by PicoSirius Red (PSR) and Collagen-1 protein, which is a major component of the extracellular matrix (ECM) surrounding the regenerating bile ducts (figure 2d). One explanation for this reduction in Collagen-1 levels is that the number of Keratin-19 ductular structures are reduced following Porcupine-i, suggesting that proliferation of these ducts is dependent on Wnt ligand (figure 2e), however we did not find this to be the case and Krt19 positivity remained the same between Porcupine-i and control groups. Similarly, the number of Desmin positive portal fibroblasts surrounding the regenerating ducts remains constant following Porcupine-i (figure 2f), indicating that inhibition of Wnt signalling results in a reduced ability of portal fibroblasts to synthesise Collagen-1, without affecting the number of ducts or fibroblasts directly.

### Myeloid Wnt5a regulates bile duct fibrosis

Based on the absence of canonical Wnt signalling in biliary regeneration we postulated that non-canonical Wnt signalling is likely driving the fibrotic phenotype which can be inhibited following Porcupine-i treatment. *Wnt4*, *Wnt5a* and *Wnt11* have all been shown to activate non-canonical Wnt signalling(17, 45, 46). Following biliary injury induced by DDC, *Wnt11* expression was low and did not change with injury (supplementary figure 3a), whereas *Wnt4* and *Wnt5a* expression increased at five days following biliary injury, when levels of p-cJUN^S73^ are high both in the murine model (supplementary figure 3a and b and figure 3a) and also in human patients with PSC (supplementary figure 4a and 4b). Moreover, during DDC induced regeneration, WNT4 is expressed exclusively by the biliary epithelium following injury, but is absent in the resting liver (supplementary figure 3c). Wnt4^KO^ mice are lethal in the first 24 hours of life due to renal defects(45), however heterozygous mice are fertile and live into adulthood. Wnt4^KO/WT^ mice were given DDC diet for five or fourteen days. Weight loss, a proxy measure for animal condition, between the Wnt4^KO/WT^ and Wnt4^WT/WT^ was the same throughout the time course studied (supplementary figure 3d). Moreover, in Wnt4^KO/WT^ compared to Wnt4^WT/WT^ littermates PSR, Collagen-1 (supplementary figure 3e) and Desmin (supplementary figure 3f) are unchanged.

**Figure 3.**
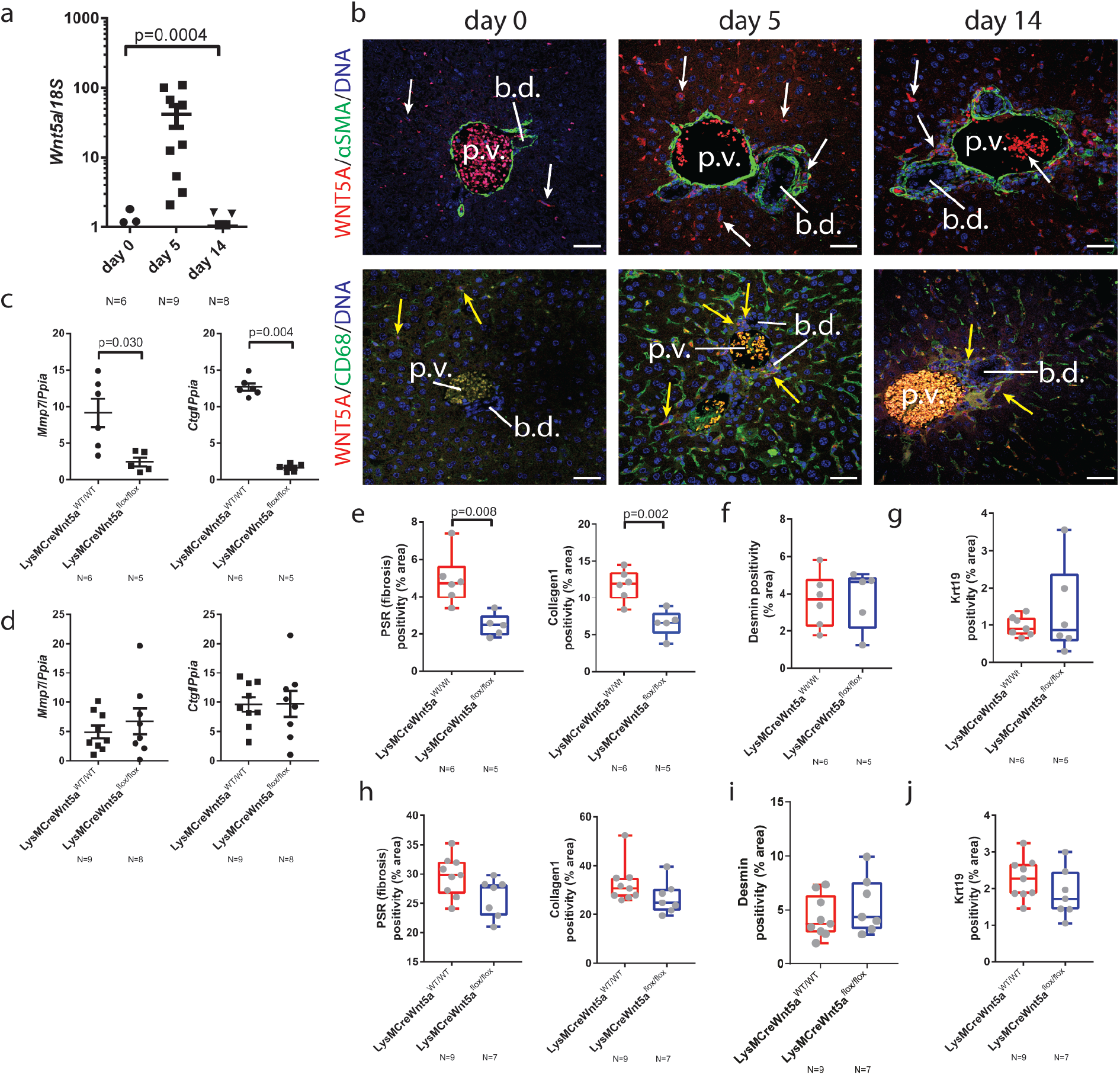
Myeloid Wnt5a regulates initiation of biliary fibrosis. (a) mRNA expression of Wnt5a from day 0, 5 and 14 DDC treated livers. (b) Immunofluorescence staining of DDC livers from day 0, 5 and 14 time points. Upper panels: Wnt5a (red), fibroblasts, αSMA (green). Lower panels: Wnt5a and Macrophages, CD68 (green). White arrows demarcate positivity and yellow arrows demarcate co-staining. (c) mRNA expression of *Mmp7* and *Ctgf* in isolated bile ducts from mice in which Wnt5a has been deleted from the myeloid lineage following by 5 days DDC or (d) 14 days DDC. (e) Image analysis of PSR and Collagen-1, (f) Desmin and (g) Keratin-19 following 5 days of DDC or (h) Image analysis of PSR and Collagen-1, (i) Desmin and (j) Keratin 19 following 14d of DDC in LysM-Cre: Wnt5aflox mice or control animals. Data are presented as mean ±SEM, statistics presented are derived from a Kruskal-Wallis test. Photomicrograph scale bars: 50μm. b.d: bile ducts, p.v. portal vein.

*Wnt5a* transcript and WNT5A protein are expressed by a number of small cells in the resting liver (figure 3b, left hand panels and supplementary figure 4c), however following DDC damage at days 5 and 14, Wnt5a positive cells become more abundant and co-localise with the macrophage marker CD68, but not the fibroblast marker αSMA (figure 3b, centre and right-hand panels). To test whether macrophage derived Wnt5a is a critical regulator of Collagen-1 deposition in bile duct repair, mice harbouring a myeloid specific deletion of Wnt5a(47, 48) (LysM-Cre/Wnt5a^flox/flox^ or LysM-Cre/Wnt5a^WT/WT^, as a control) were generated and treated with DDC for 5 or 14 days. *Mmp7* and *Ctgf* transcripts were significantly reduced in isolated bile ducts from mice without myeloid Wnt5a following five days of DDC treatment (figure 3c), however loss of these transcripts is restored by day 14 of injury (figure 3d), suggesting a compensatory mechanism controls expression of *Ctgf* and *Mmp7* later in disease. In line with this, we found that Wnt5a^flox/flox^ mice lose more weight following DDC injury than Wnt5a^WT/WT^ controls, however this difference diminishes at later time points (supplementary figure 4d). These changes in biliary gene expression were not limited to *Mmp7* and *Ctgf*, with *Dkk1* and *Wnt5b* also demonstrating myeloid Wnt5a dependant changes (supplementary figure 5e and 5f), however these changes are persistent and are not resolved at 14 days suggesting that certain transcriptional programmes still require myeloid Wnt5a for the duration of the DDC model. Similarly, a number of classical canonical and non-canonical Wnt targets failed to change following Wnt5a deletion, indicating that these are regulated through a non-myeloid Wnt5a dependant route (supplementary figure 5 a-d).

Given the ability of Porcupine-i to reduce the level of fibrosis surrounding the bile duct, mice with biliary injury with or without myeloid specific deletion of Wnt5a were analysed for levels of fibrosis. The level of PSR and Collagen-1 were significantly reduced in LysM-Cre/Wnt5a^flox/flox^ mice, compared to LysM-Cre/Wnt5a^WT/WT^ control animals, following 5 days of DDC (figure 3e). Both Desmin positive portal fibroblasts (figure 3f) and Keratin-19 positive epithelial cells (figure 3g) remained constant. By day 14 of DDC, the levels of PSR and Collagen-1 were statistically comparable between genotypes (figure 3h). Similarly, Desmin positive portal fibroblasts (figure 3i) and Keratin-19 positive biliary epithelial cells (figure 3j) remained constant, indicating that myeloid Wnt5a regulates the fibrogenic capacity of portal fibroblasts, rather than the proliferation or survival of cholangiocytes or portal fibroblasts themselves. These data suggest that Wnt5a represents an early response in biliary fibrosis that can be compensated for by other ligands later in disease or Wnt5a produced elsewhere, however, if these ligands act through common pathways then targeting non-canonical Wnt receptors would demonstrate a sustained anti-fibrotic response.

### Non-canonical Wnt receptors are expressed by biliary cells

The receptor complexes which mediate non-canonical Wnt signalling are diverse and it is unclear how they interact(15). However, a number of receptors, including Frizzled receptors and orphan receptor tyrosine kinases such as Ror1/2 and Ptk7, which themselves are known to bind Wnt5a, converge on the scaffolding protein VANGL2(18, 49, 50), (49) (schematic, figure 4a).

**Figure 4.**
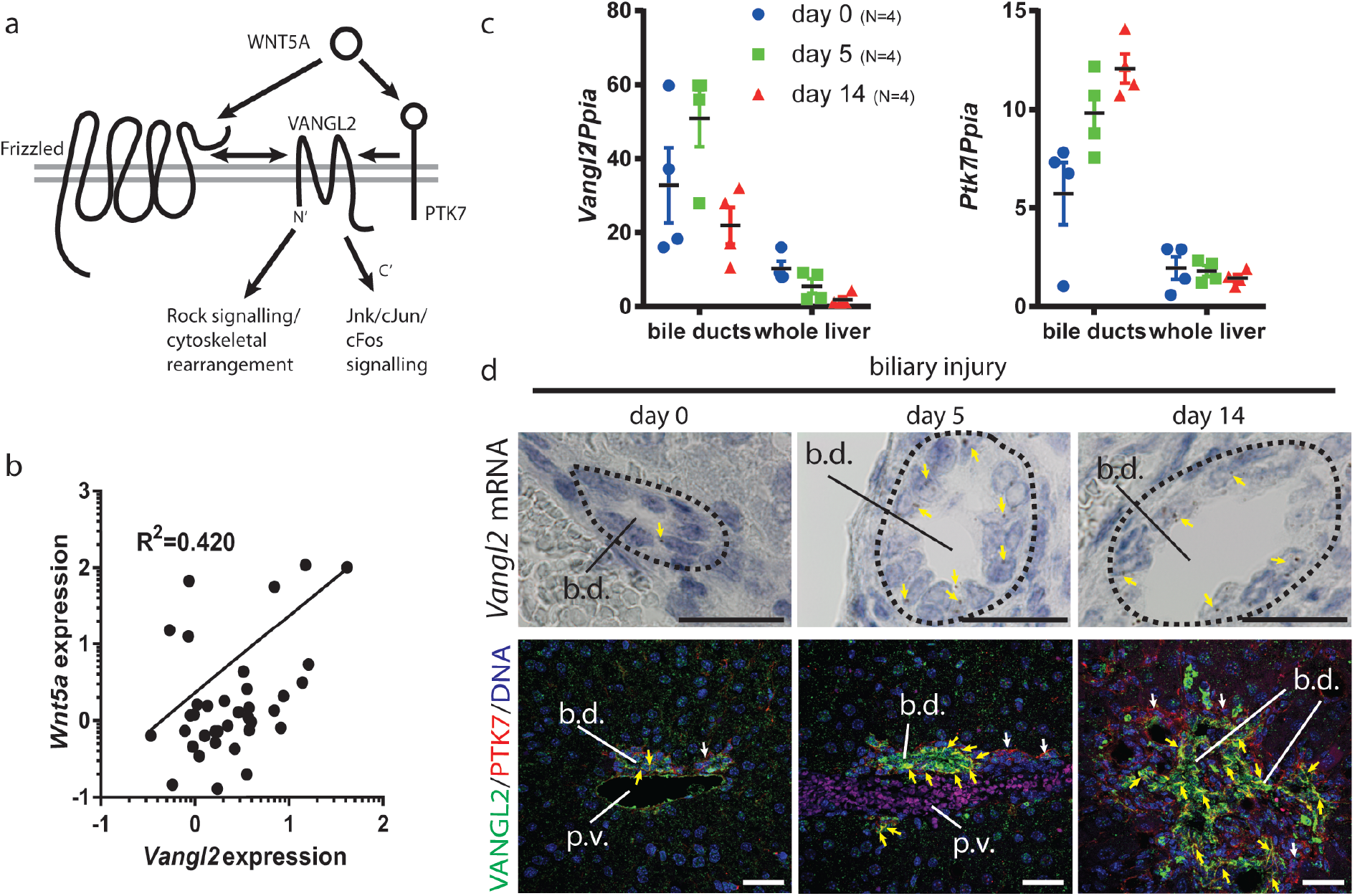
The non-canonical Wnt proteins, Vangl2 and Ptk7 are expressed in biliary regeneration. (a) A schematic demonstrating the interactions between Wnt5a, Vangl2 and Ptk7 and the non-canonical downstream pathway. (b) *Wnt5a* mRNA correlates with *Vangl2* mRNA following DDC injury. (c) *Vangl2* and *Ptk7* mRNA increases with injury in the DDC model. This increase is restricted to the bile ducts and is not seen in whole liver. (d) Upper panels: RNAScope showing that *Vangl2* mRNA is localised to the biliary epithelium, yellow arrows denote positivity. Lower panels: Immunofluorescence staining showing VANGL2 (green) and PTK7 (red) in bile ducts. White arrows denote VANGL2 and yellow arrows denote VANGL2 and PTK7 co-localisation. Data are presented as mean ±SEM, R^2^ value is derived from a linear regression analysis. Photomicrograph scale bars: 50μm. b.d: bile ducts, p.v. portal vein.

In uninjured, 5 or 14 day DDC treated mice, expression of *Wnt5a* positively correlates with that of *Vangl2* (figure 4b). Moreover, in patients with PSC, *VANGL2* mRNA levels, but not those of the homologue VANGL1, were significantly increased when compared to livers without underlying biliary disease (supplementary figure 6a). Isolated bile ducts from mice demonstrated a higher expression of *Vangl2* throughout the DDC time course than whole liver, peaking at day 5 following injury initiation. Concurrent with this, there is also an increase in expression of *Ptk7* (figure 4c), suggesting that a Vangl2-Ptk7 signalling hub is expressed in bile ducts.

Based on our previous observations, that Porcupine-i and Wnt5a deletion could regulate fibrosis, we hypothesised that Vangl2 and Ptk7 would be expressed by portal fibroblasts, which are likely to be co-isolated in our bile duct enrichment. Using RNAscope *Vangl2* mRNA could be seen rarely in biliary epithelial cells and never outside of this cellular compartment in uninjured liver. As injury progressed through to days 5 and 14, the abundance of *Vangl2* mRNA increased in biliary epithelial cells, however, no transcripts were detectable in adjacent portal fibroblasts (figure 4d, upper panels). To further validate these findings and confirm whether VANGL2 protein was indeed specifically expressed in the bile duct epithelium, liver from untreated mice or mice that had had 5 or 14 days of DDC were stained for VANGL2 and PTK7. In uninjured animals, rare VANGL2 positive cells can be seen in the bile duct epithelium, which is positive for PTK7 protein. However, at days 5 and 14 following injury there is high levels of VANGL2 and PTK7 co-expression specifically in the biliary epithelium, indicating that following injury, VANGL2 and PTK7 are both present and could function to receive non-canonical Wnt signals by biliary epithelial cells and not portal fibroblasts as hypothesised (figure 4d, lower panels and supplementary figure 6g). We confirmed that Vangl2 and its homologue Vangl1 were expressed in the biliary epithelium using a mouse line in which eGFP is fused to the C-terminus of Vangl2 (Vangl2^eGFP^ (51)). Following DDC, we detected eGFP in the membranes of biliary epithelial cells and hepatic vascular cells (particularly the large vessels rather than in the sinusoidal endothelium), but we did not see eGFP in cells adjacent to the biliary epithelium (supplementary figure 6c), eGFP also co-localises with VANGL1 protein, indicating that both Vangl2 and its interacting homologue Vangl1 are specifically expressed in biliary epithelial cells (52) (supplementary figure 6c, lower panels and supplementary figure 6d and 6e). Moreover, in tissue from patients with PSC, VANGL1 and VANGL2 localised to the biliary epithelium (supplementary figure 6b). Bile ducts isolated from the DDC time course also expressed non-canonical Wnt genes: *Dvl2* and *Dvl3* mRNA levels did not change throughout the time course (however in whole liver mRNA levels were reduced in line with increased injury). Expression of putative non-canonical Frizzled receptors (*Fzd3* and *Fzd6*) was high in isolated bile ducts compared to whole liver, but decreased throughout the time course. Finally, *Ror2*, encoding a receptor tyrosine kinase that is known to interact with PTK7, is transcriptionally increased in bile ducts following 5 days of injury (supplementary figure 6e). Notably, PTK7 protein could also be detected on portal fibroblasts, however there was no Vangl2 expressed in these cells and therefore the role of PTK7 here is unclear (supplementary 6g, lower panels). We could not rule out that PTK7 directly receives the Wnt5a signal, thereby conferring the fibrogenic phenotype we have described in the absence of Vangl2.

To determine whether Vangl2-Ptk7 signalling can account for the changes in fibrosis seen following Porcupine-i or myeloid Wnt5a deletion, bile duct organoids were generated from mice containing either Vangl2^flox/flox^ (with loxP sites spanning the transmembrane domains of Vangl2, figure 5a(53)) or Vangl2^WT/WT^ and were infected with lentivirus expressing Cre from the CMV promoter. Following Cre expression, VANGL2 protein and *Vangl2* mRNA expression are lost (denoted as Vangl2^ΔTM^, figure 5b and 5c), whilst *Vangl1* mRNA remains unchanged (figure 5c). Vangl2 functions upstream of Jnk/cJUN signalling in embryonic development(18) and in breast cancer cells(26). We have shown that Porcupine-i alters Jnk/cJun in biliary epithelial cells, therefore we sought to determine whether this phenotype was mediated through Vangl2.

**Figure 5.**
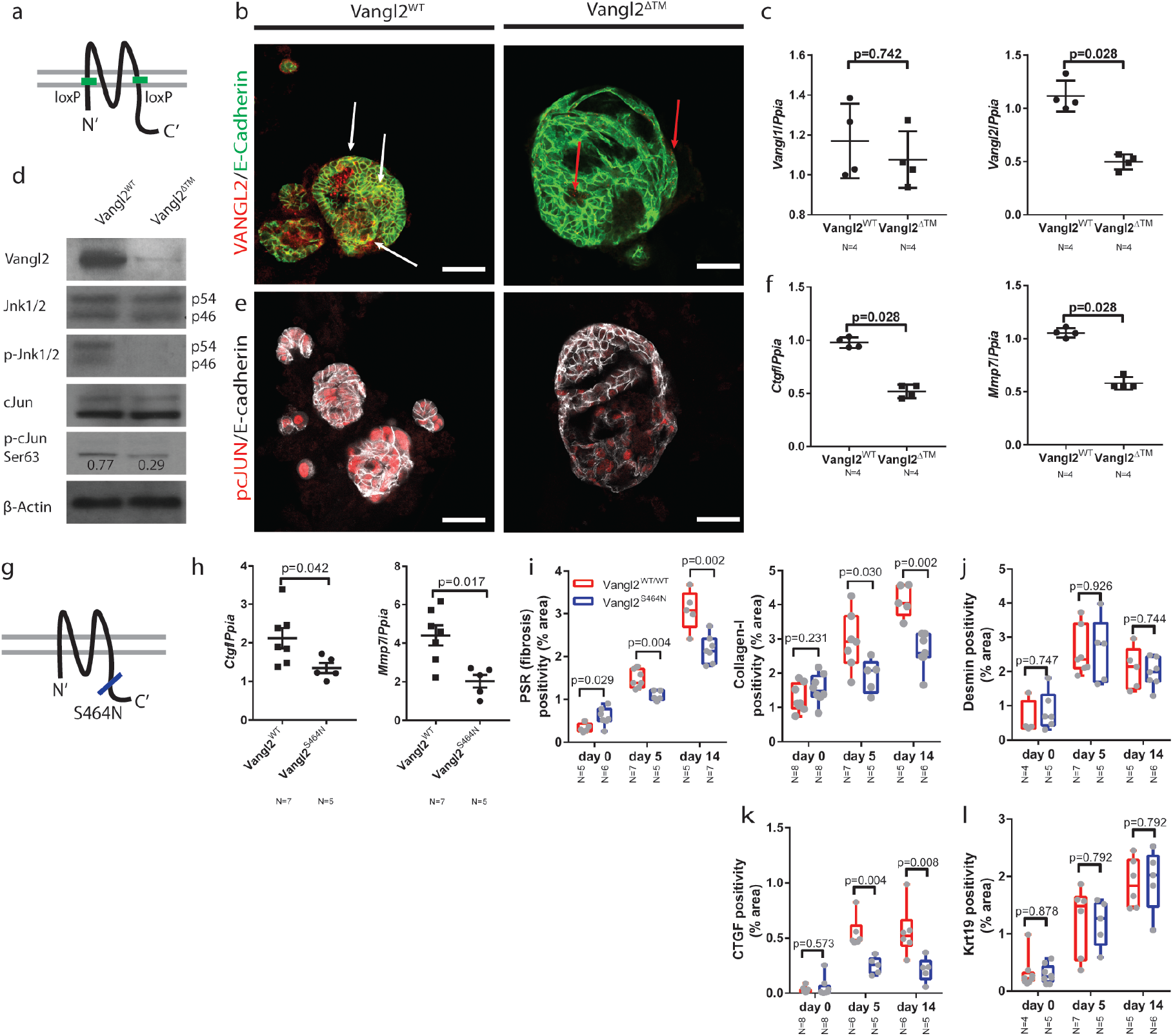
Vangl2 regulates non-canonical Wnt signalling in the adult bile duct. (a) Schematic representing the location of the Vangl2^loxP^ sites in the Vangl2^flox/flox^ mouse line. (b) Immunocytochemistry for VANGL2 (red) and E-cadherin (green) in CMV-Cre:Vangl2^flox/flox^ or CMV-Cre-Vangl2^flox/flox^ organoids. White arrows denote co-positivity, red arrows denote red puncta which are not localised to the membrane in the organoids that have lost Vangl2 (c) mRNA expression of Vangl1 and Vangl2 following recombination of Vangl2^flox^ in bile duct organoids (d) Western blots showing VANGL2, DVLl2, JNK1/2, pJNK1/2^T183/Y185^, cJUN, p-cJUN^S63^ and β-actin levels in organoids derived from Vangl2^flox^ or Vangl2^WT^ mice. p-cJUN^S63^ is normalised to total cJUN (e) pcJUN^S73^ (red) and E-Cadherin (white) immunocytochemistry on organoids derived from Vangl2^flox/flox^ or Vangl2^WT^ mice and infected with Lv-CMV-Cre. (f) mRNA expression of *Ctgf*, *Mmp7*, *Mmp14* and *Cd44* in organoids derived from Vangl2^flox^ or Vangl2^WT^ mice. (g) Schematic showing the location of the Vangl2^S464N^, Looptail mutation. (h) mRNA expression of *Ctgf* and *Mmp7* from isolated bile ducts derived from mice given 5 days DDC. (i) PSR and Collagen-1 staining from mice carrying a single copy of the Vangl2^S464N^ mutation compared to those with two WT alleles. (j) Desmin positivity from mice carrying a single copy of the Vangl2^S464N^ mutation compared to those with two WT alleles. (k) CTGF positivity from mice carrying a single copy of the Vangl2^S464N^ mutation compared to those with two WT alleles (l) Krt19 positivity from mice carrying a single copy of the Vangl2^S464N^ mutation compared to those with two WT alleles Data are presented as mean ±SEM, statistics presented are derived from a Mann-Whitney test in the cases where two groups are compared. Photomicrograph scale bars: 50μm.

In the Vangl2^WT^ organoids, levels of total-JNK1/2 remain constant, however, following deletion of Vangl2, levels of p-JNK^T183/Y185^ are reduced. Similarly, whilst total cJUN levels remain constant in Vangl2^WT^ organoids, the level of p-cJUN^S63^ is reduced by 2.6-fold (figure 5d) in Vangl2^ΔTM^ organoids. Moreover, this reduction in p-cJUN was confirmed immunocytochemically by staining for p-cJUN^S73^ in Vangl2^ΔTM^ organoids compared to bile duct organoids in which Vangl2 is intact (figure 5e). Bile duct organoids are solely comprised of epithelial cells and can be used to ask questions about the downstream transcriptional changes following Vangl2 alteration without the confounding effects of other cell types found around the damaged bile duct. mRNA for both *Ctgf* and *Mmp7* was significantly down regulated in Vangl2^ΔTM^ organoids compared to control organoids (figure 5f), similarly *Mmp14* and *Cd44* mRNA was reduced, however *Sparc* and *Mmp3* mRNA was increased following Vangl2 deletion (supplementary figure 7a).

Mutations in the C-terminal domain of Vangl2 are known to affect its distribution and function i*n vivo* and have been shown to reduce levels of downstream signalling(54, 55). Vangl2^S464N^ mutant mice (also known as *Looptail*, figure 5g) were given biliary injury for up to 14 days. Following five days of injury, Vangl2^S464N^ mice had reduced mRNA expression of *Ctgf* and *Mmp7* in isolated bile ducts (figure 5h). Moreover, following injury, Vangl2^S464N^ (56) mutant mice lost significantly more weight following injury than their Vangl2^WT^ littermates (supplementary figure 7b). Livers from uninjured, five day and 14 day DDC mice were stained for PSR, Collagen-1 and Desmin. Vangl2^S464N^ mutant mice have a higher resting level of PSR and Collagen-1 than their Vangl2^WT^ littermates. However, following damage, the levels of PSR and Collagen-1 deposited in the Vangl2^WT^ are significantly higher than the levels in Vangl2^S464N^ mutants (figure 5i). Moreover, the levels of Desmin or Krt19 positive cells do not change between mutant and wild-type animals (figure 5j and 5k), however levels of CTGF do (figure 5l), suggesting that there is an alteration in the capacity of portal fibroblasts to synthesise Collagen-1, rather than a change in the number of fibrogenic cells *per se*.

Vangl2^S464N^ mice have developmental abnormalities in a number of tissues and it is possible that the phenotype here is due to a developmental phenotype that is continued to adulthood. To overcome this, Krt19CreER^T^ mice(57), in which Cre is specifically activated in biliary epithelial cells, were crossed with Vangl2^flox/flox^ mice(53) to generate K19CreER^T^/Vangl2^flox/flox^, K19CreER^T^/Vangl2^flox/WT^ or K19CreER^T^/Vangl2^WT/WT^ mice (figure 6a). These mice also harbour a R26R^LSLtdTomato^ allowing cells in which Cre has been activated to be tracked using RFP. Following tamoxifen administration (after which mice are denoted as Vangl2^ΔTM^), mice were given DDC for up to fourteen days. Following five days of DDC, RFP positive cell numbers remains constant across all genotypes, demonstrating that biliary epithelial loss of Vangl2 is not cytotoxic *in vivo* (supplementary figure 7c). As expected, in Vangl2^ΔTM^ mice, RFP positive cells lose Vangl2 staining in the membrane (supplementary figure 7d). Five days following activation of K19CreER^T^, p-JNK^T183/Y185^/RFP dual positive cells can be found in Vangl2^WT/WT^ mice, however, in mice that have a Vangl2^ΔTM^ allele, RFP positive cells are largely negative for p-JNK^T183/Y185^, placing Vangl2 upstream of JNK activation (figure 6b, left-hand panels). Similarly, in RFP positive cells with intact Vangl2, p-cJUN^S73^ can be seen in the nucleus, however, in RFP cells from tamoxifen treated mice, Vangl2^ΔTM^ cells do not express p-cJUN^S73^ whilst adjacent, non-RFP cells remain positive (figure 6b, right-hand panels). In tandem with decreased phosphorylation of JNK and cJUN, we also found a significant decrease in the expression of CTGF throughout the DDC time course (figure 6c) in K19CreER^T^/Vangl2^ΔTM^ and K19CreER^T^/Vangl2^ΔTM/WT^ mice compared to K19CreER^T^/Vangl2^WT/WT^ controls. In line with this, K19CreER^T^Vangl2^ΔTM^ and K19CreER^T^Vangl2 ^ΔTM/WT^ mice also have significant reduction in the levels of PSR and Collagen-1 at five days and fourteen days following injury initiation (figure 6d and 6f) and improved body weights (supplementary figure 7e), without changes in the number of Desmin positive fibroblasts (figure 6e and 6g) when compared to K19CreER^T^Vangl2^WT/WT^ controls.

**Figure 6.**
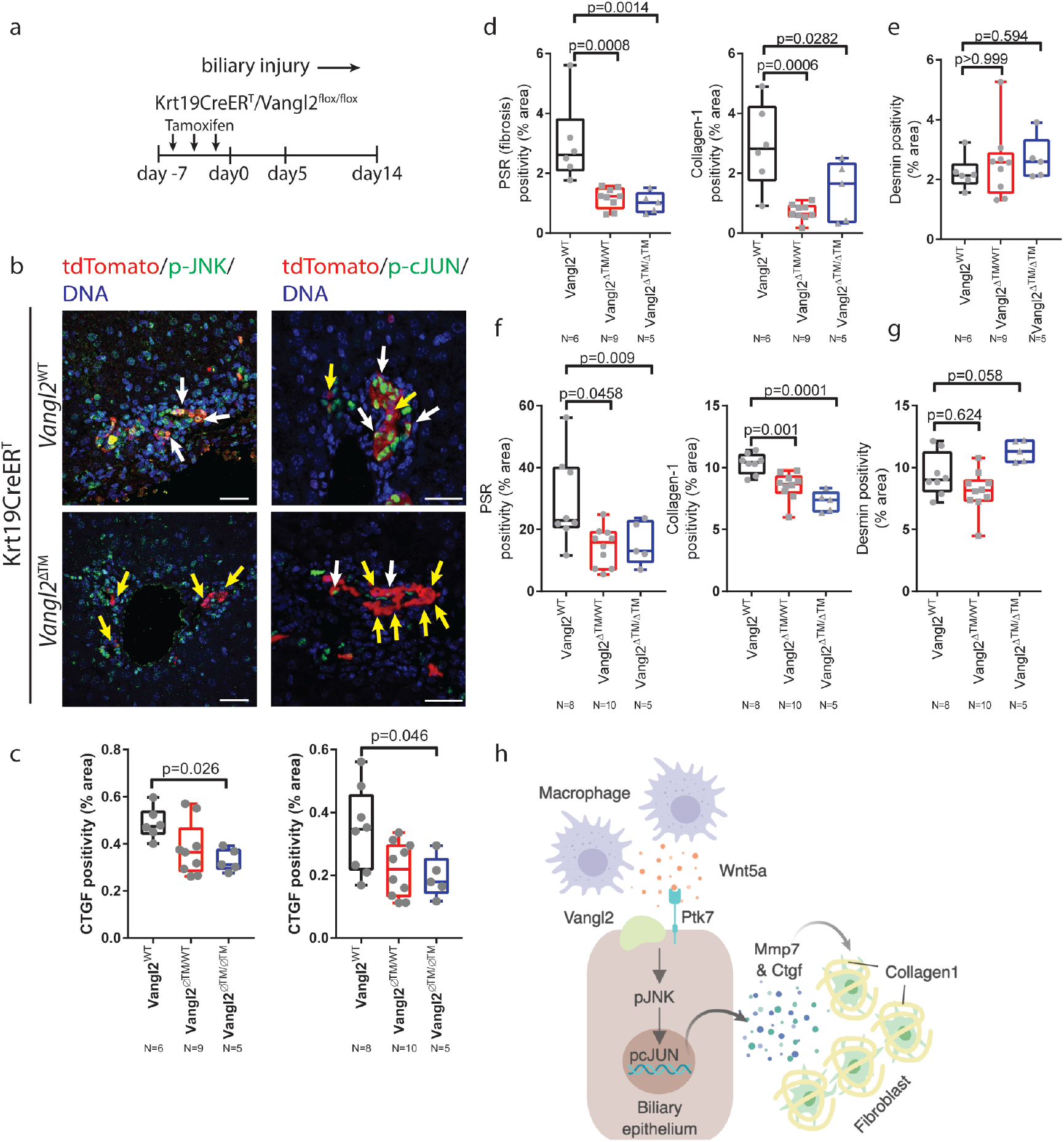
Deletion of Vangl2 in the biliary epithelium regulates biliary fibrosis in vivo. (a) Schematic of Vangl2 deletion in K19CreER^T^Vangl2^flox/flox^ or control mice undergoing DDC injury. Tamoxifen is administered prior to initiation of DDC induced damage. Mice also contain a R26R^tdTomato^ to identify cells in which Vangl2 has been recombined. (b) Immunofluorescence showing tdTomato positivity (red), denoting cells in which Cre is activated. p-JNK1/2^T183/Y185^ (left-hand panels) and p-cJUN^S73^ (right-hand panels) in K19CreER^T^Vangl2^flox/flox^ or Vangl2^WT^ mice. White arrows denote tdTomato positive cells that are also positive for pJNK or pcJUN, Yellow arrows denote tdTomato positive cells in which pJNK or pcJUN is absent. (c) Quantification of CTGF staining in K19CreER^T^Vangl2^ΔTM^, Vangl2^WT/ΔTM^ or Vangl2^WT/WT^ (d) PSR and Collagen 1 staining and (e) Desmin staining 5 days following DDC initiation or (f) PSR and Collagen 1 and (g) Desmin staining 14 days following DDC initiation in K19CreER^T^ mice with Vangl2^WT^, Vangl2^flox/WT^ or Vangl2^flox/flox^. (h) model of the signalling network described in this study. Data are presented as mean ±SEM, statistics presented are derived from a Kruskal-Wallis test. Photomicrograph scale bars: 50μm.

Together, our data indicate that non-canonical Wnt signalling is important for the initiation of the fibrotic response in biliary injury. This is achieved through the local recruitment of Wnt5a expressing myeloid cells, which then activate JNK/cJUN signalling in biliary epithelial cells. The consequence of this activation is the transcriptional activation of *Mmp7* and *Ctgf*, which despite being expressed by biliary epithelial cells themselves, co-localises with adjacent fibrogenic cells indicating that factors expressed by the biliary epithelium directly influence the fibrogenic capacity of adjacent fibroblasts, and thereby regulate Collagen-1 deposition and scarring (figure 6h).

## Discussion

The understanding of Wnt signalling in adult tissues and diseases is dominated by the canonical, β-catenin Wnt signalling pathway, which is known to drive proliferation of cells in multiple developmental, disease and cancer contexts. Studies unpicking non-canonical Wnt signalling pathways have lagged behind and it is unclear whether these signalling pathways, which are critical for normal development, are also required for adult physiology and disease

Here, we have demonstrated that in the adult liver, a number of central, non-canonical signalling components are expressed by inflammatory macrophages and biliary epithelial cells. In development these components have been implicated in maintaining tissue patterning and stabilising architectural changes, whilst in the adult these signalling components are implicated in the early establishment of a fibrotic microenvironment. In our model and in correlative human samples Wnt5a expressed by myeloid cells is received by a Vangl2/Ptk7 mediated complex which, through control of Mmp7 and Ctgf regulates the *de novo* deposition of Collagen-1 in a non-cell autonomous way. In this way, non-canonical Wnt signalling acts to integrate a number of cell types to facilitate biliary scarring. Whilst excessive scarring is detrimental to liver function, deposition of scar tissue arises due to fundamental insufficiencies in epithelial regeneration. Whilst the stromal cells surrounding the regenerating bile duct produce scar, they are also known to provide a number of ligands that stimulate epithelial regeneration including Jagged-1 (5, 58) and Hedgehog (59, 60). Therefore, having a mechanism though which the epithelium can regulate the stroma adjacent to it allowing for sufficient co-evolution of the regenerative microenvironment, whilst regulating scar formation, must be a centrally important aspect of tissue repair.

Two previous reports (32, 42) have demonstrated differing effects of myeloid Wnts in mouse models of cholestatic damage. Our data, deleting Wnt5a from myeloid cells, appears to support the conclusions drawn by Okabe et al, who showed that loss of myeloid Wnt production through the loss of *Wls* resulted in a reduced fibrotic response in DDC model of biliary injury. Furthermore, our data support their speculation, that it is likely to be a Wnt dependant, but β-catenin independent pathway that regulates this biliary repair process *in vivo*. However, our data does contradict these reports in some aspects, as we fail to see changes in the number of bile ducts formed following *Wnt5a* deletion. This could be explained by the Wnt ligands, with diverse functions, expressed by macrophages and would explain the *Wls* phenotype seen by Okabe et al (34, 61–63). Similarly, the differences we see between our study and a more chronic TAA model, in which *Wls* is deleted, can be explained by parenchymal damage secondary to chronic biliary disease(64). It has previously been shown that Wnt signalling is needed for hepatocyte regeneration(9, 65, 66) following injury, an increase in fibrosis would be expected if loss of myeloid Wnts results in a failure to drive this epithelial repair process in the parenchyma.

Modulating the canonical Wnt signalling pathway has proven difficult in the non-cancerous context, potentially due to the relatively limited amount of Wnt activity that is required in homeostasis(67, 68). Moreover, off-target effects of canonical Wnt inhibitors have limited the usefulness of this pathway therapeutically. However, recent developments have identified well-tolerated canonical Wnt pathway inhibitors that are being used in a range of oncological indications. We suggest that in the liver, the non-canonical Wnt signalling pathway is activated in biliary disease and that this pathway can be modulated to minimise biliary scarring while maintaining the cellular complexity of the regenerating microenvironment. Given that in many biliary diseases injury occurs in iterative and cyclical patterns, understanding the initiating signals and targeting these could lead to significant patient improvement.

## Materials

### Mouse models

All mice were maintained in 12h light/dark cycles and had access to food and water ad libitum in accordance with UK Home Office Regulations. All experiments were performed under license PPL: 70/8150 held by Dr Luke Boulter, mice were euthanised by an escalating dose of CO_2_.

For the DDC mouse model. 6-8-week-old, male CD1 mice were fed 0.1% 5-diethoxycarbonyl-1,4-dihydrocollidine (DDC) in their diet for up to 14 days and provided with normal drinking water. Mice which drop >20% of their bodyweight were given DDC food softened with water. In studies were mice received the Porcupine inhibitor, LGK974 experimental mice were dosed with 5mg/Kg LGK974, twice daily via oral gavage (the vehicle for this was 0.5% methylcellulose and 0.5% Tween-80 in dH2O). Mice were dosed for 48h prior to the administration of DDC to ensure inhibited Wnt ligand production prior to injury. Effectiveness of LGK974 was confirmed through reduction of GS staining adjacent to central veins and is known to be Wnt ligand dependent.

Vangl2^eGFP^ were provided by Dr Ping Chen, Emory University, USA and were maintained on a CD1 background. Looptail mice (Vangl2^S464N^, Harwell, UK and provided by Dr Charlotte Dean) were maintained on a C3H background and throughout these studies, wild-type litter mates served as a control. Krt19CreER^T^:R26RtdTomato:Vangl2^f/f^ mice were generated by crossing the Krt19CreER^T^ mouse line initially with the tdTomato Ai14 line from Jax labs. Mice Heterozygous for the K19CreER^T^ knock-in and homozygous for tdTomato were then crossed with Vangl2flox mice, kindly provided by Prof Deborah Henderson, Newcastle. In this study, all Krt19CreER^T^:R26RtdTomato:Vangl2^f/f^ mice were positive for tdTomato and CreER^T^ and were then either homozygous, heterozygous or wild-type for the Vangl2flox allele. Induction of Cre was achieved through repeat administration of Tamoxifen (three times for 1 week) at a concentration of 4mg per mouse for each injection. Tamoxifen was diluted in 10% molecular grade ethanol: 90% corn oil. LysMCre:Wnt5a^f/f^ mice were generated by crossing the LysMCre line kindly provided by Dr Steven Jenkins, University of Edinburgh, with Wnt5a^flox/flox^ mice purchased from Jax labs. Mice were maintained as Cre positive and either homozygous, heterozygous or wild-type for the floxed allele. All transgenic mice were genotyped by Transnetyx Inc.

## Human tissue

### RNA isolation and RT-PCR

RNA was isolated from both tissue and organoids using a modified Trizol Method. Briefly, tissue was homogenized in Trizol and RNA collected from the aqueous phase following chloroform extraction. RNA was precipitated from the aqueous phase using isopropanol. Precipitated RNA was applied to an RNeasy Mini Spin Column (Qiagen) and the manufacturer’s protocol was followed. RNA was suspended in RNAse/DNAse free H2O and quantified using a Nanodrop (Thermofisher). 1ug total RNA was treated with gDNA wipe-out buffer (Qiagen) and reverse transcribed using the Quantitect reverse transcription kit (Qiagen) as per the manufacturer’s instructions. For quantitative PCR 10ug total cDNA was used per reaction and was run using FastStart SYBR Green reagent (Roche), cycle conditions were defined by the manufactures protocol. All qRT-PCR was run using a Lightcycler 480-II. All primer sets used in this study were purchased from Qiagen (Table 1)

**Table 1.** qPCR primers used in this study.

| Table 1 – qPCR primers used in this study. |  |  |
| --- | --- | --- |
|  | Provider | Catalogue Number |
| **Mouse** |  |  |
| axin2 | Qiagen | QT00126539 |
| bmp4 | Qiagen | QT00111174 |
| ccnd1 | Qiagen | QT00154595 |
| cd44 | Qiagen | QT00173404 |
| ctgf | Qiagen | QT00096131 |
| cyr61 | Qiagen | QT00245217 |
| dcn1 | Qiagen | QT00131068 |
| dkk1 | Qiagen | QT00146307 |
| dvl2 | Qiagen | QT00112154 |
| dvl3 | Qiagen | QT01079092 |
| egfr | Qiagen | QT00101584 |
| fzd3 | Qiagen | QT00147917 |
| fzd6 | Qiagen | QT00109998 |
| gas1 | Qiagen | QT00253190 |
| igfbp1 | Qiagen | QT00114716 |
| lef1 | Qiagen | QT00148834 |
| mmp14 | Qiagen | QT01064308 |
| mmp3 | Qiagen | QT00107751 |
| mmp7 | Qiagen | QT00110012 |
| myc | Qiagen | QT00096194 |
| pkcθ | Qiagen | QT00171325 |
| ptk7 | Qiagen | QT00144032 |
| rybp | Qiagen | QT00310275 |
| sparc | Qiagen | QT00161721 |
| stat6 | Qiagen | QT01042503 |
| tpm | Qiagen | QT00137354 |
| vangl1 | Qiagen | QT00494235 |
| vangl2 | Qiagen | QT00128422 |
| wnt11 | Qiagen | QT00103663 |
| wnt4 | Qiagen | QT00104622 |
| wnt5a | Qiagen | QT00164500 |
| wnt5b | Qiagen | QT00169708 |
| **Human** |  |  |
| VANGL1 | SAbiosciences | PPH21116A |
| VANGL2 | SAbiosciences | PPH12377B |
| WNT5A | SAbiosciences | PPH02410A |
| **Rat** |  |  |
| cd44 | SAbiosciences | PPR42408A |
| ctgf | SAbiosciences | PPR46426B |
| mmp14 | SAbiosciences | PPR44815A |
| mmp3 | SAbiosciences | PPR48487B |
| mmp7 | SAbiosciences | PPR44767C |
| ptk7 | SAbiosciences | PPR52923C |
| sparc | SAbiosciences | PPR53743A |
| vangl2 | SAbiosciences | PPR54102B |
| wnt5a | SAbiosciences | PPR50278A |

### Tissue fixation and Immunohistochemistry

Following euthanasia, livers were flushed through with 20ml physiological saline. Dissected liver lobes were then fixed overnight in 10% neutral buffered formalin in PBS. Following fixation, tissue was dehydrated in 70% ethanol and processed to wax using a tissue processor. 4µM tissue sections were cut and rehydrated through graduated alcohols. Tissue sections were antigen retrieved as per Table 2. For DAB staining of antigens, tissue sections were treated with 10% H_2_O_2_ washed and then native streptavidin and biotin was blocked using a commercial avidin and biotin blocking kit. Following this, sections were blocked with a universal protein block. Tissue sections were incubated over-night with primary antibodies diluted in antibody diluent. Following washing, biotin conjugated secondary antibodies were incubated with tissue sections as per Table 2. Sections were further incubated with ABC vectorstain and colour was developed using DAB substrate. For tyramide stained sections, the avidin/biotin step is omitted and HRP conjugated secondary antibodies and Alexa488 or Alexa594 conjugated tyramide was used as per the manufacturer’s instructions. For directly conjugated immunofluorescent antibodies, the same protocol was performed, omitting the H_2_O_2_ and Avidin/Biotin blocking stage.

**Table 2.**
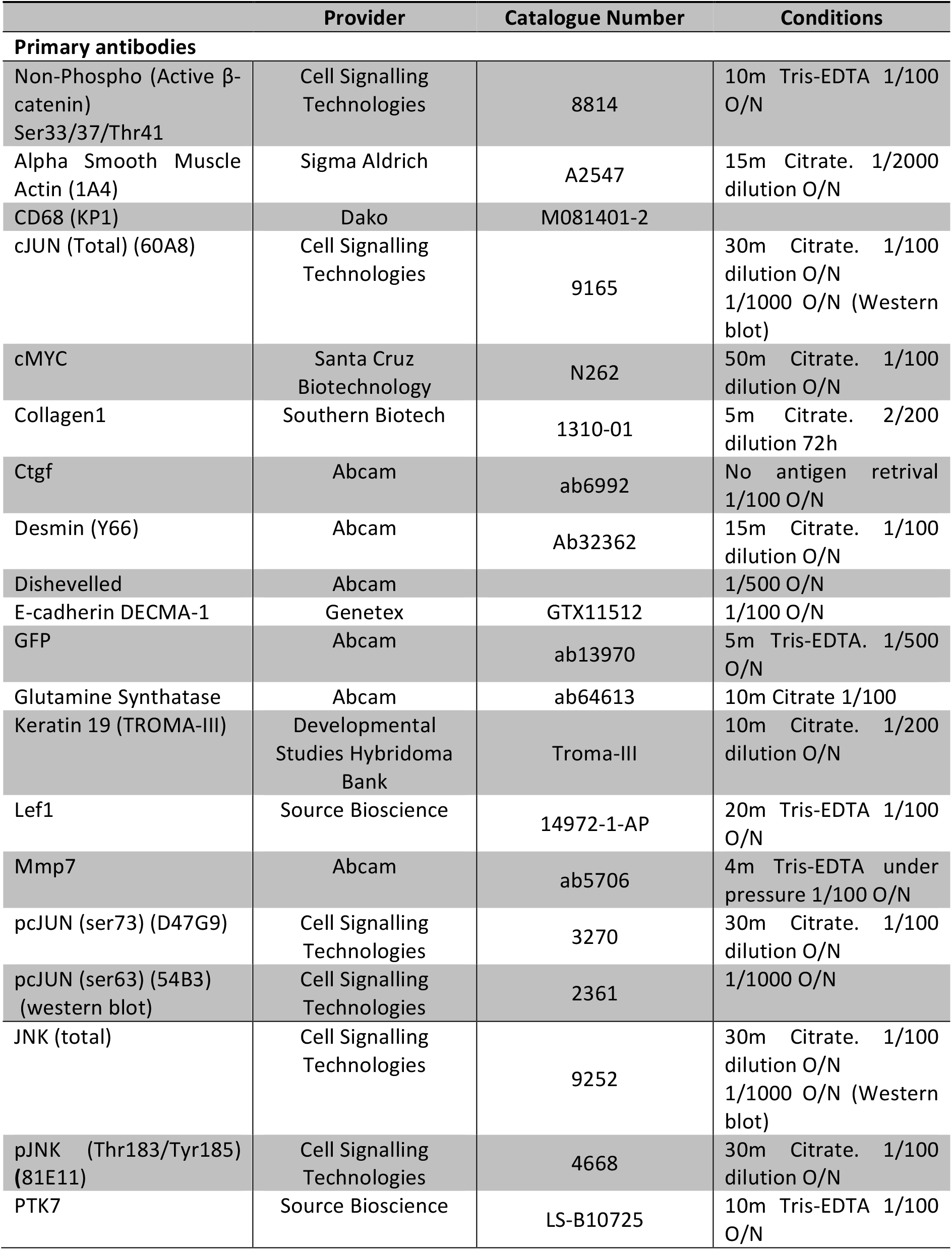
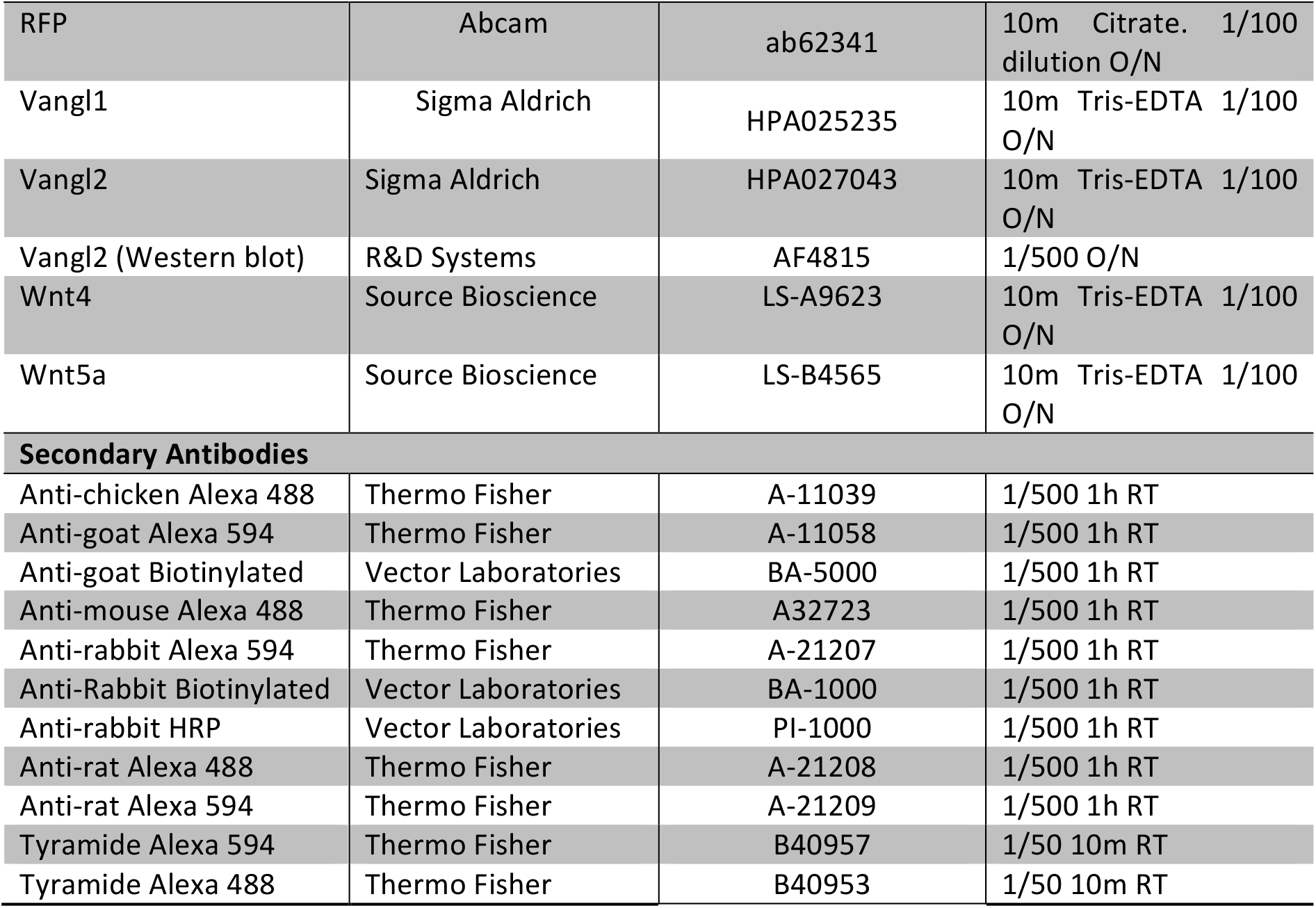
Antibodies used in this study.

DAB stained sections were imaged on either an Olympus Dotslide, Hamamatzu Nanozoomer or an Olympus BX53 upright microscope. Immunofluorescence was imaged using a Nikon A1R confocal microscope.

### RNAScope

RNAscope was performed on formalin fixed liver tissue. 4µm sections were used throughout and all RNAScope performed in this study was done by Aquila Histoplex, Edinburgh.

### Bile duct isolation

To isolate bile ducts from both uninjured and injured livers dissected liver was chopped into 5mm^3^ pieces and digested in DMEM/F12 media containing Collagenase-IV (Roche) and DNASe-I (Roche). Following digestion and dissociation, bile ducts become obvious as parenchyma is digested away. Bile ducts are strained through a 70µm filter and extensively washed in PBS to remove any residual cells. Bile ducts are then used for downstream applications.

### Organoid culture

Liver organoids were derived from isolated bile ducts. Briefly, isolated bile ducts were resuspended in 100% GFR Matrigel. Within 24h bile ducts form closed structures and within 48h budding can be seen from the duct. Following expansion, these ducts are removed from Matrigel by incubating with ice-cold Versene. Organoids are dissociated with pipetting and then re-plated in fresh 100% Matrigel. This process is repeated to expand organoids. The growth media used in this study consists of a base media of DMEM/F12 supplemented with Glutamax, Penicillin/Streptomycin, Fungizone and HEPES. Just prior to feeding, the base media is supplemented with HGF, EGF, FGF10, Gastrin, Nicotinamide, N-Acetylcystine, B-27, Forskolin, Y-27632 (ROCK inhibitor), A83-01 (TGFβ inhibitor) and Chir99021 (GSK3β inhibitor).

To evaluate the effects of Vangl2 knockout on bile duct organoids (Vangl2^flox/flox^) versus control organoids (Vangl2^WT/WT^) Cre was expressed from the CMV promotor, *in vitro*, by infection of organoid structures with Lv-CMV-Cre (University of Edinburgh, SuRF facility) at an MOI of 5. For infection media was supplemented with 1/100 of Transdux reagent. For assay win which signalling was monitored in response to genotype growth media was withdrawn from organoid cultures and organoids were cultured in base media containing B-27, Forskolin, Nicotinamide and N-Acetylcystine alone for 48h, at which point organoids were lysed for analysis.

### Western blotting

Isolated bile ducts or organoids were lysed in RIPA buffer containing phosphatase (ThermoFisher) and protease (MiniComplete, Roche) inhibitors. Protein quantification was determined using Pierce BCA reagent (Pierce) and quantified using a nanodrop. The standard curve for protein quantification was derived from the BCA reagent handbook using Albumin standards provided. 20ug total protein was loaded onto a 4-12% NuPage Bis-Tris gel (ThermoFisher). Prior to running, proteins were reduced with NuPage LDS sample buffer (4x) and NuPage Sample Reducing Agent (10x). All gels were run using NuPage MOPS SDS Running buffer containing NuPage Antioxident. Proteins were transferred onto PVDF membrane (Amersham) using NuPage Transfer Buffer. Following transfer, membranes were blocked in either 5% BSA (Sigma Aldrich) in TBST or 5% dried milk (Marvel) in TBST. Membranes were incubated with primary antibodies (Table 2) at 4°C overnight. Following washing with TBST membranes were incubated with HRP-conjugated secondary antibodies (Table 2) at room temperature for 1h. Following washing signal was developed using ECL (Pierce).

### Image analysis

Image analysis was conducted using ImageJ and macros written by Dr Tim J Kendall. Macros are available on request.

### Statistics

All experimental groups were analysed for normality using a D’Agostino-Pearson Omnibus test. Groups that were normally distributed were compared with either a two-tailed student’s t-test (for analysis of two groups) or using one way ANOVA to compare multiple groups. Non-parametric data was analysed using a Wilcoxon-Mann Whitney U test when comparing two groups or a Kruskall-Wallis test when comparing multiple non-parametric data. Throughout p<0.05 was considered significant. Data is represented at mean with S.E.M for parametric data or median with S.D. for non-parametric data.

### Study Approval

All animal experiments were approved by the University of Edinburgh local ethics committee and were licensed by the UK Home Office. All patient material contained in this manuscript was approved by the NHS Lothian Bio resource ethics committee. No prospective tissue was collected in this study.

## Author Contributions

DW performed experiments and analysis, RM, EJ and AR performed experiments, PC provided advice and the Vangl2^GFP^ knock-in mouse line through collaboration. CD collaborated on provision of the Vangl2^LP^ mouse line and Vangl2^LP^ tissues. DH generated and provided the Vangl2flox line. TJK provided pathological support, wrote the macros for image analysis, contributed to the direction of the project and edited the manuscript. LB conceived the project, provided funding for the project, conducted experiments and analysed data, compiled and wrote the manuscript and led the project.

## Acknowledgements

The authors would like to thank Guoqiang Gu for originally providing the Krt19CreER^T^ mouse line and Ed Boulter-Comer for technical reading of the manuscript. This work was funded by Primary Sclerosing Cholangitis (PSC) Support, The Alan Moremont Memorial Foundation (AMMF, The Cholangiocarcinoma Charity) and core funding provided to the MRC Human Genetics Unit, by the Medical Research Council.

## Notes

Conflict of Interest Statement: The authors have declared that no conflict of interest exists.

