## Supplementary Materials for "Non-canonical Wnt signalling initiates scarring in biliary disease"

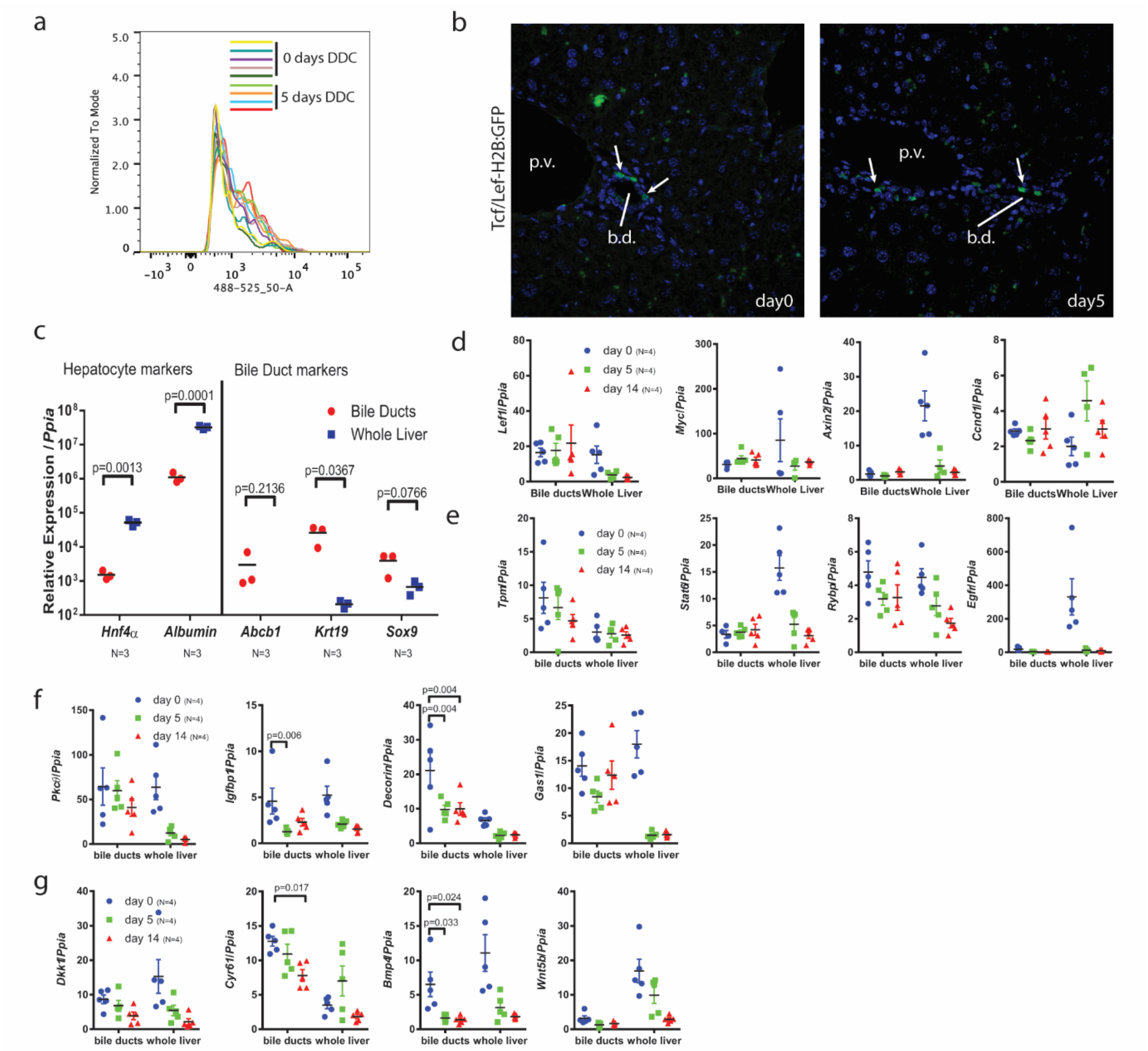


**Supplementary Figure 1. Canonical, β-catenin dependant, Wnt signalling is not regulated in biliary disease.** (a) Flow cytometry from isolated biliary cells from TCF/LEF-H2B: GFP mutant mice showing levels of GFP in biliary cells in resting liver and day 5 DDC injury. (b) Immunohistochemistry from TCF/LEF-H2B: GFP mice either untreated or following 5 days of DDC. White arrows denote GFP positive biliary cells. (c) mRNA for bile duct and hepatocyte markers in whole liver and isolated bile ducts as a demonstration of biliary enrichment. (d-g) mRNA expression of canonical Wnt signalling (*Lef1*, *Myc*, *Axin2*, *Ccnd1*), cFOS/cJUN (*Tpm*, *Stat6*, *Rybp*, *Egfr*, *PKCϑ*, *Igfbp1*, *Dcn1*, *Gas1*) and YAP/Taz (*Dkk1*, *Cyr61*, *Bmp4*, *Wnt5b*) target genes in whole liver and isolated bile ducts following 0, 5 and 14 days of DDC injury. Data are presented as mean ±SEM, statistics presented are derived from a Kruskal-Wallis test. N=3 per group for C and N=4 per group for all other analysis.

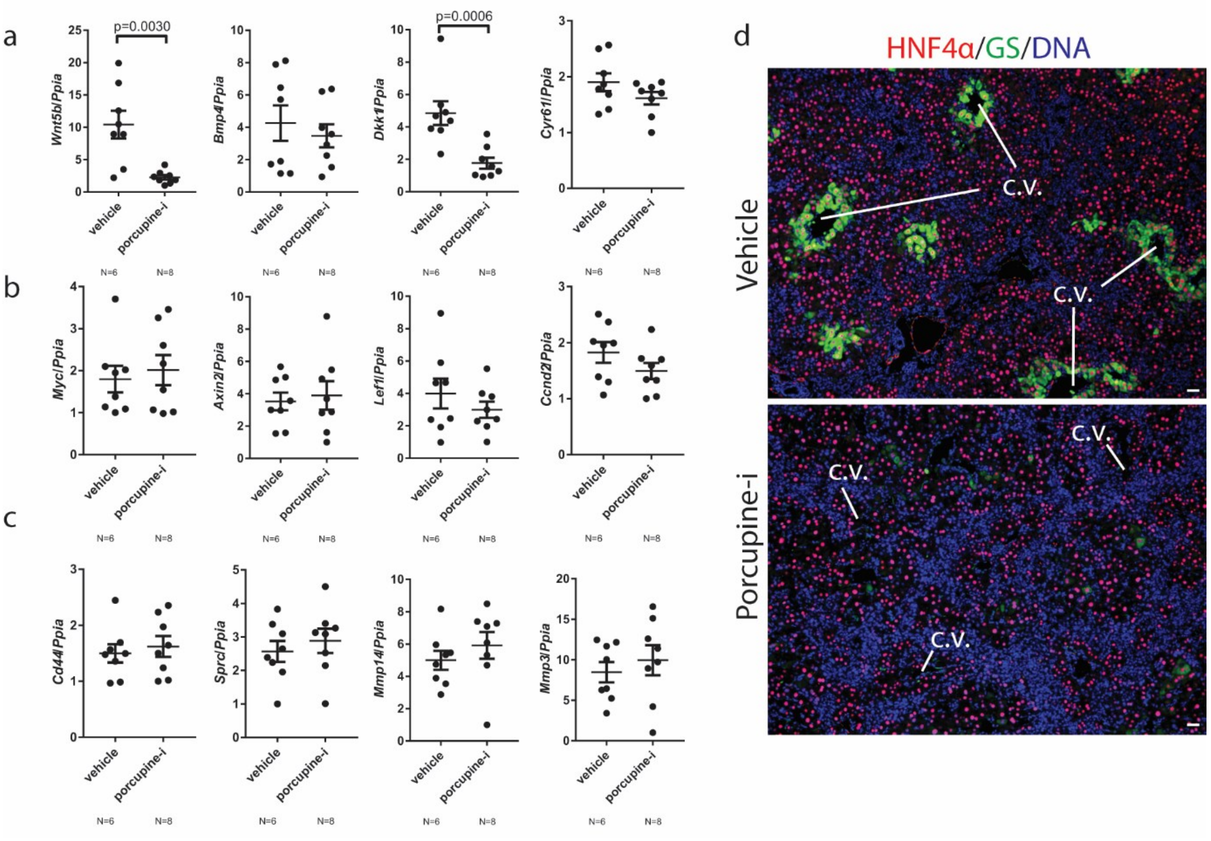


**Supplementary Figure 2. Porcupine inhibition alters targets of non-canonical Wnt signalling, but not canonical Wnt signalling in the biliary epithelium.** (a) mRNA expression of *Wnt5b*, *Bmp4*, *Dkk1* and *Cyr61* in isolated bile ducts from mice following five days DDC injury treated with Porcupine inhibitor or vehicle. (b) Canonical Wnt target genes, *Myc*, *Axin2*, *Lef1* and *Ccnd1* in isolated bile ducts from mice following five days DDC injury treated with Porcupine inhibitor or vehicle (c) cJUN/cFOS targets *Cd44*, *Sprc*, *Mmp14* and *Mmp3* in isolated bile ducts from mice following five days DDC injury treated with Porcupine inhibitor or vehicle. (d) Immunofluorescence staining showing HNF4α (red), Glutamine Synthase (green) in vehicle and Porcupine inhibitor treated liver. Data are presented as mean ±SEM, statistics presented are derived from a Mann-Whitney test. c.v. denotes central veins.

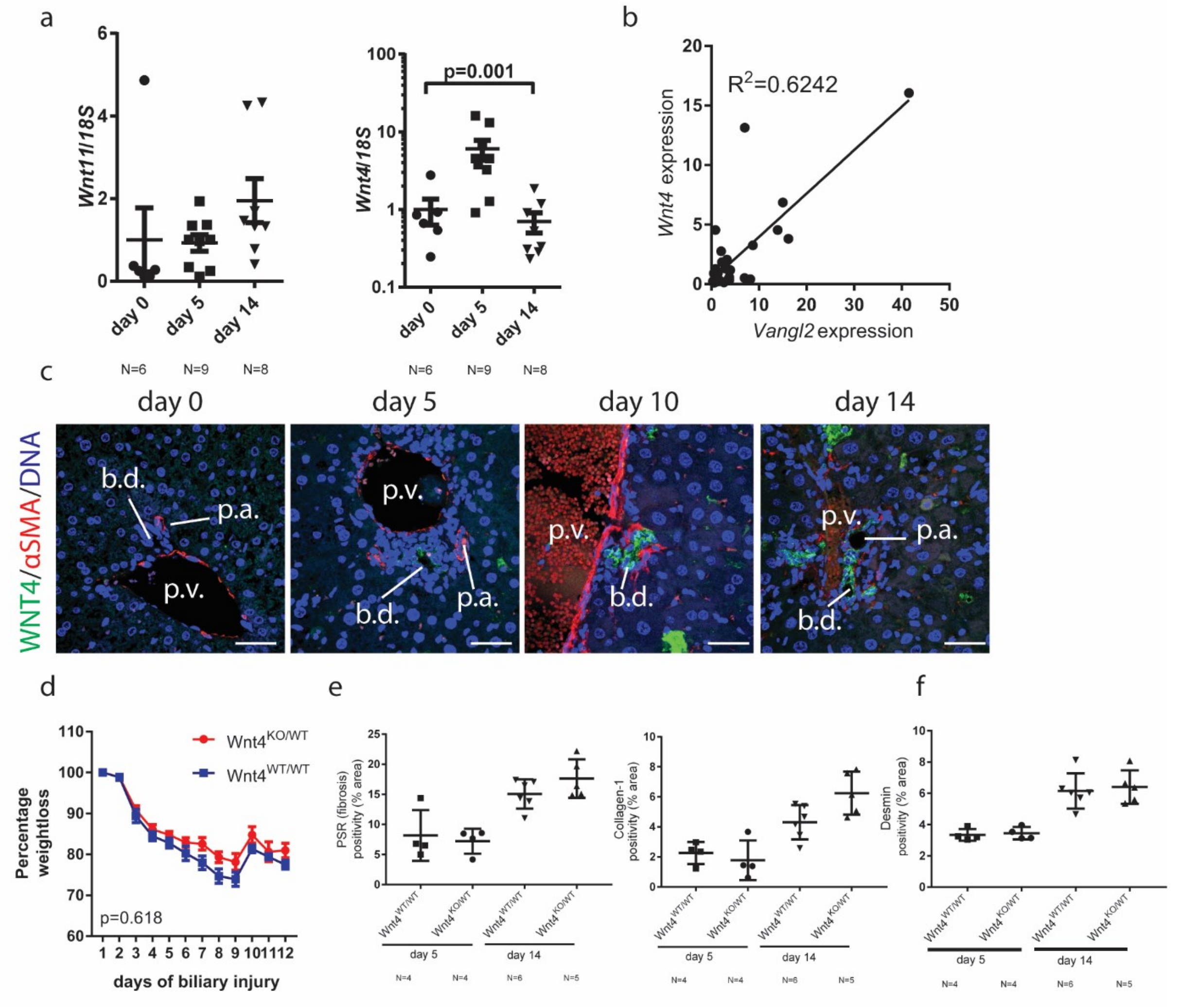


**Supplementary Figure 3. Wnt4 haploinsufficicency does not regulate ECM deposition in biliary injury.** (a) mRNA expression of *Wnt11* and *Wnt4* throughout the DDC time course at days 0, 5 and 14. (b) Wnt4 and Vangl2 mRNA correlation over the DDC time course. (c) Immunohistochemistry of Wnt4 and fibroblasts throughout the DDC timecourse at d0, d5 and d14. (d) Weightloss in $Wnt4^{KO/WT}$ and $Wnt4^{WT/WT}$ mice over the time course of DDC injury. (e) Quantification of PSR and Collagen-1 immunostaining in $Wnt4^{KO/WT}$ and $Wnt4^{WT/WT}$ livers treated with 5 or 14 days of DDC. (f) Quantification of Desmin immunostaining in $Wnt4^{KO/WT}$ and $Wnt4^{WT/WT}$ livers treated with 5 and 14 days of DDC. Data are presented as mean ±SEM, statistics presented are derived from a Mann-Whitney test when two groups are being compared or a Kruskal-Wallis test when > 2 groups are being compared. $R^2$ is calculated by a linear regression model. Comparisons between $Wnt4^{WT}$ and $WNT4^{KO/WT}$ were derived using a 2-way ANOVA. Photomicrograph scale bars: 50µm. b.d. denotes bile ducts, p.a. portal arteries and p.v. portal veins.

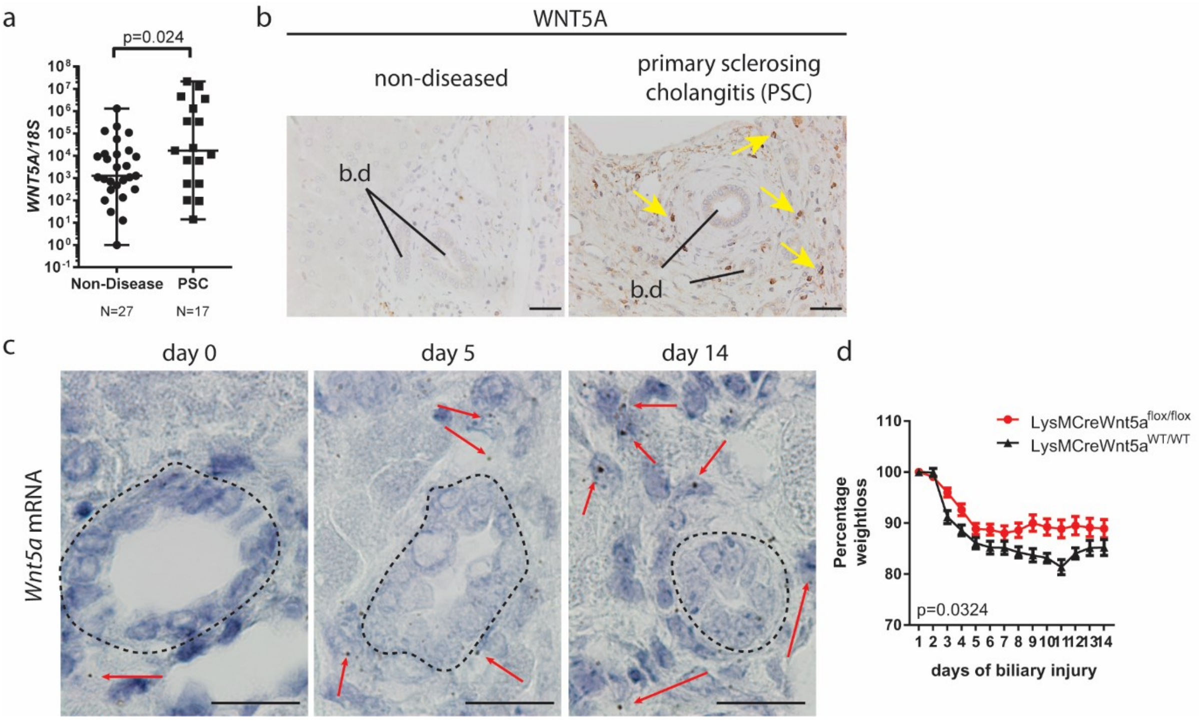


**Supplementary Figure 4. Wnt5a is expressed in biliary injury.** (a) mRNA expression of *WNT5A* in patients with no liver disease or with primary sclerosing cholangitis. (b) WNT5A protein expression (brown) in non-diseased human liver compared to liver from a patient with primary sclerosing cholangitis. Yellow arrows denote positivity (c) RNAscope, showing *WNT5A* mRNA throughout the DDC time course. Red arrows denote positivity. (d) Weight loss of LysMCreWnt5a$^{WT/WT}$ and LysMCreWnt5a$^{flox/flox}$ mice with DDC injured liver throughout the DDC experimental time course. Photomicrograph scale bars: 50µm. b.d. denotes bile ducts, p.a. portal arteries. Data are presented as mean ±SEM, statistics presented are derived from a Mann-Whitney test. Statistics comparing LysM-Cre:Wnt5a$^{flox/flox}$ vs Wnt5a$^{WT/WT}$ mice were derived from a two way ANOVA.

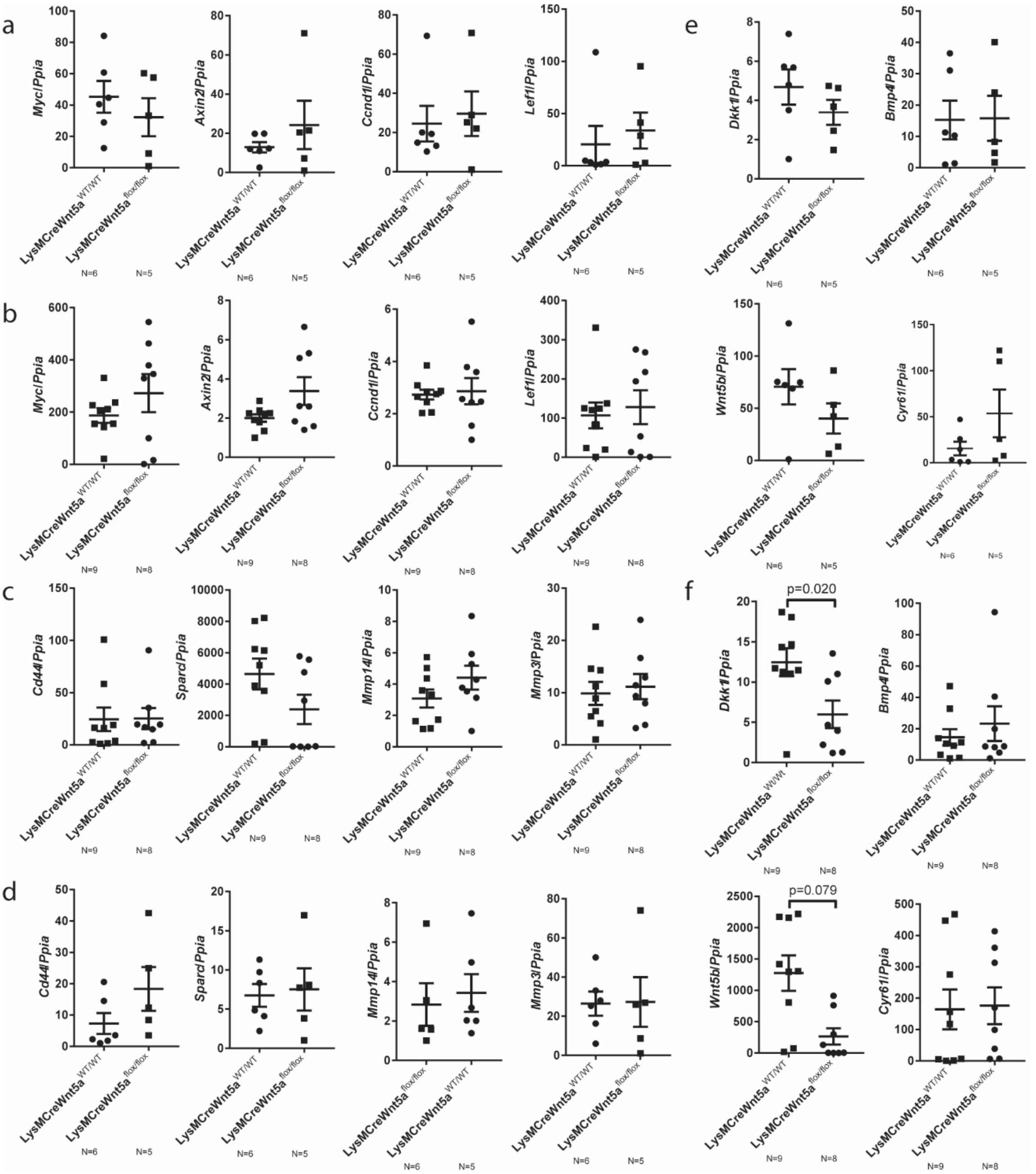


**Supplementary Figure 5. Wnt pathway targets following deletion of Myeloid Wnt5a.** (a) mRNA expression of canonical Wnt signalling targets (*Myc*, *Axin2*, *Ccnd1*, *Lef1*) in isolated bile ducts from LysMCreWnt5a$^{flox/flox}$ versus LysMCreWnt5a$^{WT/WT}$ 5 days following DDC injury. (b) mRNA expression of canonical Wnt signalling targets (*Myc*, *Axin2*, *Ccnd1*, *Lef1*) in isolated bile ducts from LysMCreWnt5a$^{flox/flox}$ versus LysMCreWnt5a$^{WT/WT}$ 14 days following DDC injury. (c) mRNA expression of cJun/cFos targets (*Cd44*, *Sparc*, *Mmp14*, *Mmp3*) in isolated bile ducts from LysMCreWnt5a$^{flox/flox}$ versus LysMCreWnt5a$^{WT/WT}$ 5 days following DDC injury. (d) mRNA expression of cJun/cFos targets (*Cd44*, *Sparc*, *Mmp14*, *Mmp3*) in isolated bile ducts from LysMCreWnt5a$^{flox/flox}$ versus LysMCreWnt5a$^{WT/WT}$ 14 days following DDC injury. (e) mRNA expression of Yap/Taz targets (*Dkk1*, *Bmp4*, *Wnt5b*, *Cyr61*) in isolated bile ducts from LysMCreWnt5a$^{flox/flox}$ versus LysMCreWnt5a$^{WT/WT}$ 5 days following DDC injury. (f) mRNA expression of Yap/Taz targets (*Dkk1*, *Bmp4*, *Wnt5b*, *Cyr61*) in isolated bile ducts from LysMCreWnt5a$^{flox/flox}$ versus LysMCreWnt5a$^{WT/WT}$ 14 days following DDC injury. Data are presented as mean ±SEM, statistics presented are derived from a Mann Whitney test.

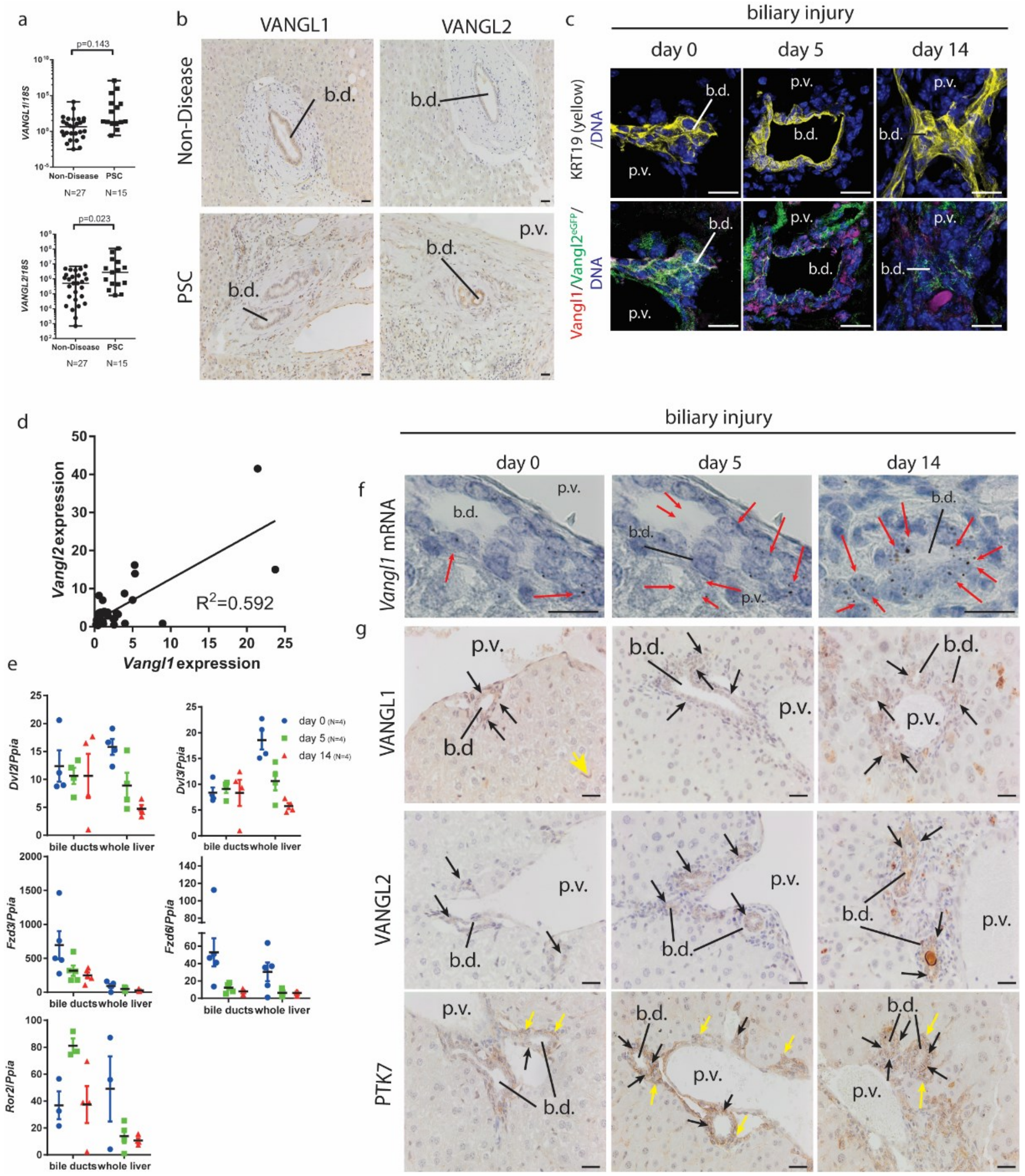


**Supplementary Figure 6 – Biliary epithelial cells express non-canonical Wnt receptors.** (a) mRNA expression of *VANGL1* and *VANGL2* in non-diseased liver compared to patients with primary sclerosing cholangitis. (b) VANGL1 and VANGL2 protein expression in non-diseased human compared to patients with primary sclerosing cholangitis. (c) Upper panels, Keratin-19 (yellow) immunoflourescence denoting bile ducts at day 0, day 5 and day 14 of DDC injury in Vangl2$^{eGFP}$ knock-in mice. Lower panels, VANGL1 (red) and GFP (Vangl2$^{eGFP}$) immunofluorescence throughout the DDC time course. (d) Correlation of *Vangl1* and *Vangl2* mRNA levels in the DDC model of biliary injury. (e) mRNA expression of *Dvl2* and *Dvl3* and putative non-canonical Frizzled receptors *Fzd3* and *Fzd6* as well as non-canonical receptor tyrosine kinases *Ror1* and *Ror2* in isolated bile ducts and whole liver form mice given DDC for 0, 5 and 14 days. (f) RNAscope for *Vangl1* mRNA in ducts from the DDC model at day 0, day 5 and day 14. Red arrows denote positivity. (g) Immunohistochemistry for VANGL1, VANGL2 and PTK7 protein in DDC livers from day 0, day 5 and day 14 of injury, black arrows denote biliary positivity, yellow arrows denote positivity in non-biliary cells. Data are presented as mean ±SEM, statistics presented are derived from a Mann Whitney test or $R^2$ is derived from a regression test. Photomicrograph scale bars: 50µm. b.d: bile ducts, p.v. portal vein.

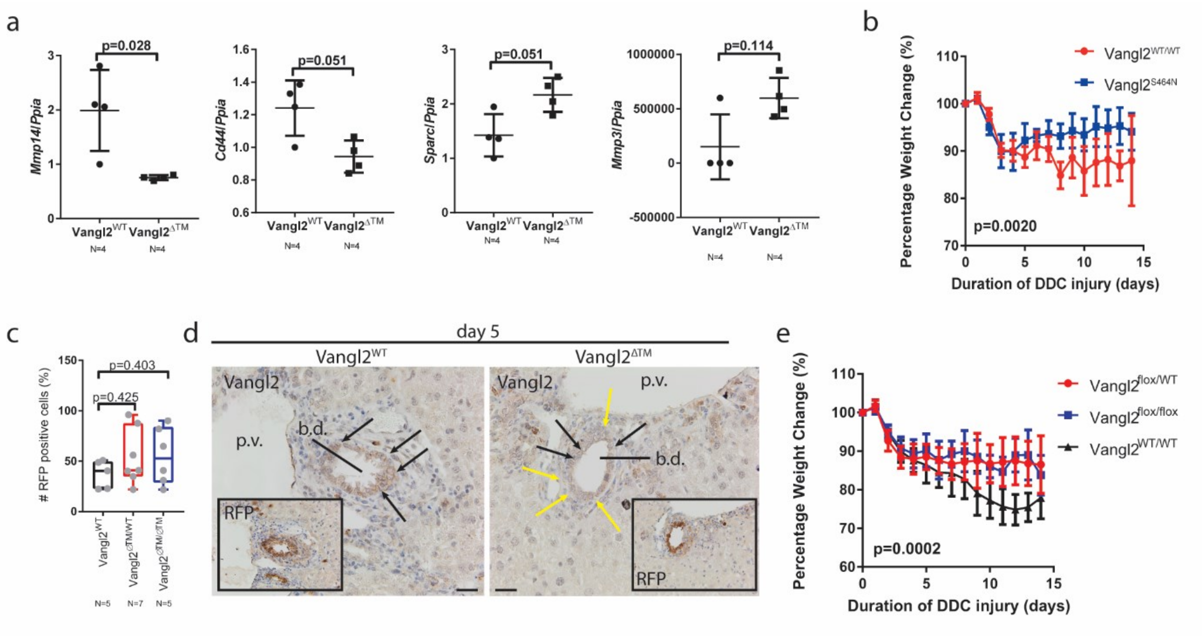


**Supplementary Figure 7 – Hypomorphic mutations or knockout of Vangl2 result in an altered response to biliary damage.** (a) mRNA expression of *Mmp14*, *Cd44*, *Sparc* and *Mmp3* in vangl2$^{\Delta TM}$ organoids compared to Vangl2$^{WT}$ controls. (b) Body weights of Vangl2$^{WT}$ compared to Vangl2$^{S464N}$ mice over the DDC timecourse. (c) Image analysis of RFP positivity after five days DDC in Krt19CreER$^{T}$ mice with either Vangl2$^{\Delta TM}$, Vangl2$^{\Delta TM/WT}$ or Vangl2$^{WT/WT}$. (d) Photomicrogrpahs of serial section of Krt19CreER$^{T}$/Vangl2$^{WT}$ versus Krt19CreER$^{T}$/Vangl2$^{\Delta TM}$ livers following five days DDC, black arrows demarcate RFP positive, Vangl2 positive cells, Yellow arrows RFP positive Vangl2 negative cells. (e) Krt19CreER$^{T}$ mice with either Vangl2$^{\Delta TM}$, Vangl2$^{\Delta TM/WT}$ or Vangl2$^{WT/WT}$ body weights following DDC administration. Photomicrograph scale bars: 50µm. b.d. denotes bile ducts, p.v. portal vein. Data are presented as mean ±SEM, statistics presented are derived from a Mann-Whitney test. Statistics comparing weight loss were derived from a two way ANOVA.
